# Analysis of ageing-dependent thiol oxidation reveals early oxidation of proteins involved in core proteostasis functions

**DOI:** 10.1101/2023.05.08.539783

**Authors:** Katarzyna Jonak, Ida Suppanz, Julian Bender, Agnieszka Chacinska, Bettina Warscheid, Ulrike Topf

## Abstract

Oxidants have a profound impact on biological systems in physiology and under pathological conditions. Oxidative post-translational modifications of protein thiols are well-recognized as a readily occurring alteration of proteins. Changes in protein thiol redox state can modify the function of proteins and thus can control cellular processes. However, chronic oxidative stress causes oxidative damage to proteins with detrimental consequences for cellular function and organismal health. The development of techniques enabling the site-specific and quantitative assessment of protein thiol oxidation on a proteome-wide scale significantly expanded the number of known oxidation-sensitive protein thiols. However, lacking behind are large-scale data on the redox state of proteins during ageing, a physiological process accompanied by increased levels of endogenous oxidants. Here, we present the landscape of protein thiol oxidation in chronologically aged wild-type *Saccharomyces cerevisiae* in a time-dependent manner. Our data determine early oxidation targets in key biological processes governing the *de novo* production of proteins, folding, and protein degradation. Comparison to existing datasets reveals evolutionary conservation of early oxidation targets. To facilitate accessibility and cross-species comparison of the experimental data obtained, we created the OxiAge Database, a free online tool for the research community that integrates current datasets on thiol redoxomes in aged yeast, nematode *Caenorhabditis elegans,* fruit fly *Drosophila melanogaster*, and mouse *Mus musculus*. The database can be accessed through an interactive web application at http://oxiage.ibb.waw.pl.

## Introduction

Proteins are essential for a large variety of functions in an organism. They provide the building blocks of the cell, the basis for cell signalling, metabolic activity, transport of molecules, DNA replication, structure, response to stimuli, and many more activities that are crucial for normal cellular function. Thus, proteins must be protected from damage to confer their cellular function. Intracellular oxidants produced during cellular stress or ageing pose an imminent threat to proteins, lipids, and DNA in the cell (Cooke et al., 2003; Mesika & Reichmann, 2019; Pizzino et al., 2017). High or chronic levels of reactive oxygen species (ROS) are often linked with pathological conditions (Krisko & Radman, 2019). During cellular ageing increasing endogenous levels of ROS are thought to accelerate the ageing process. In contrast, low levels of ROS were established as messengers in young and proliferating cells. Here, specific oxidation-sensitive amino acid residues become reversibly oxidized to change the function of a given protein temporarily. Thiol groups in the protein cysteine (Cys) residues are known to be reactive and can become readily oxidized (Paulsen & Carroll, 2013). Reversible thiol modifications linked with changes in the function or activity of proteins are commonly referred to as redox switches (Antelmann & Helmann, 2011; Groitl & Jakob, 2014). Many proteome-wide studies in various species mapped oxidation-sensitive cysteine residues, but only for a minority the biological function of the redox switch is known (Araki et al., 2016; Gould et al., 2015; McDonagh et al., 2014; Meng et al., 2021; Topf et al., 2018; van der Reest et al., 2018; Xiao et al., 2020). Redoxome studies commonly found that cellular proteins are largely present in a reduced state under non-stressed physiological conditions (Brandes et al., 2013; Go et al., 2011; Meng et al., 2021). Some oxidative thiol modifications are reversed through the action of specific cellular reduction systems that control the proteins’ redox state (Calabrese et al., 2017; Morgan et al., 2013; Rhee, 2016; Ulrich & Jakob, 2019; Winterbourn & Hampton, 2015). Over-oxidation of thiols damages the proteins irreversibly and can make them more prone to degradation or aggregation (Paulsen & Carroll, 2013), a phenomenon also observed during biological ageing (David, 2012; Finelli, 2020; Hartl, 2017; Hipp et al., 2019; Reeg & Grune, 2015). While the cause and consequences of protein damage during ageing have a prevalent place in research, too little attention was paid to the advantages of initially increasing ROS levels during ageing, mediating reversible protein thiol modifications and potential adaptation of stress responses. Thus far, only a handful of studies have addressed the proteome-wide changes in protein redox state during eukaryotic ageing. Studies on chronologically aged yeast *S. cerevisiae*, strain DBY746 (Brandes et al., 2013), and aged nematode *C. elegans* (Knoefler et al., 2012) revealed a general trend of increased proteome oxidation during ageing. More recent work performed in mice (Xiao et al., 2020) and fruit flies (Menger et al., 2015), argues that the selectivity of the oxidation targets but not the global changes in proteome oxidation might be a cellular strategy to regulate the response to ageing at the later stages of life. The study of Xiao *et al*. contributes a comprehensive database of more than 9,400 proteins reversibly oxidized in young and old mice, aged four and twenty months, respectively. However, due to the lack of data for the intermediate stages of ageing, it is difficult to address the redox changes of the proteome during the early phase of adult life. Data on thiol oxidation at the time of early ageing might help to resolve the conundrum of the role of reversible oxidation as one of the first signals for the cell to fight the ageing process. Although the previous studies in *S. cerevisiae* (Brandes et al., 2013), *C. elegans* (Knoefler et al., 2012), and *D. melanogaster* (Menger et al., 2015) identified early ageing-dependent changes in protein oxidation, the small number of quantified thiol-containing peptides (∼300 for yeast and fruit fly, ∼150 for worm throughout entire time course) do not allow visualization of the full temporal redox landscape in an ageing eukaryotic system.

In this study, we provide a large dataset on the *S. cerevisiae* thiol protein oxidation landscape during chronological ageing. It is assumed that restriction of cell proliferation is one of the main causes of age-related accumulation of macromolecular defects, leading to a deterioration of tissues and organ failures (Longo et al., 2012). Therefore, chronological ageing in *S. cerevisiae* is a model system of non-dividing cells, which is commonly utilized to better understand the mechanisms of human ageing. We found that key processes of the protein homeostasis network are targeted for thiol oxidation during the early stages of ageing. We refer to these phenomena as “early oxidation”. Proteins involved in translation, *de novo* folding and the ubiquitin-proteasome system appeared to be rapidly oxidized before bulk oxidation of the proteome accompanied by cell death occurs. A cross-species comparison of the aged redoxomes showed that similar age-dependent oxidation targets can be found in mice. We integrate our yeast data and the published work in other species and provide the results in the form of a database named “OxiAge” openly accessible to the research community (http://oxiage.ibb.waw.pl). The database will enable in-depth studies on the function of evolutionarily conserved cysteine residue oxidation during organismal ageing.

## Results

### Analysis of the redox state of yeast proteome during chronological ageing

To investigate the regulation of cellular response to ageing through reversible protein thiol oxidation, we performed a time-resolved, quantitative redoxome study in chronologically aged *S. cerevisiae* (Figure 1A). We used the established OxICAT method (Leichert et al., 2008) to determine in the *in vivo* conditions the oxidation status of redox-sensitive protein thiols via quantitative mass spectrometry (MS). Samples were collected at the mid-logarithmic growth phase (Log) and during the stationary phase of chronologically aged yeast (Figures 1A and B). The experimental design enabled to monitor the reversible oxidation of protein thiols at specific phases of ageing: in young proliferating cells (Log, mid-logarithmic phase), just after diauxic shift (day 0), during early ageing (day 3), mid-point of ageing (day 6), and late stage of ageing (day 9). A total of 9,213 unique cysteine-containing peptides (Cys-peptides) in 3,139 proteins (out of 5,478 cysteine-containing proteins described in the yeast proteome, UniProtKB; see the Methods section) were detected in three biological replicates per time point (Supplementary Table 1). Due to the low accuracy of the third biological replicate of day 3 and an unproportionally high number of unquantified peptides in comparison to other replicates, we removed this replicate from further downstream analysis. To increase the robustness of our dataset, we retained only Cys-peptides quantified in at least two biological replicates of each time point. The final filtered dataset contains a total of 3,178 quantified thiol-containing peptides, matching 3,564 cysteine sites on 1,772 proteins (Supplementary Table 2). Among the quantified Cys-peptides, 2,822 contain a single cysteine residue, 324 peptides two, and 32 peptides at least three residues (Supplementary Figure 1A). Following we will refer to the filtered dataset as the “yeast OxiAge” dataset (Supplementary Table 2). The quantified redox changes were highly reproducible between biological replicates of each time point with average Pearson’s correlation coefficients of 0.93 for the dataset containing only non-imputed values and the dataset containing additionally imputed missing values (Supplementary Figure 1B). Principle-component analysis (PCA) showed that biological replicates were separated between the different time points (Supplementary Figure 1C).

**Figure 1.**
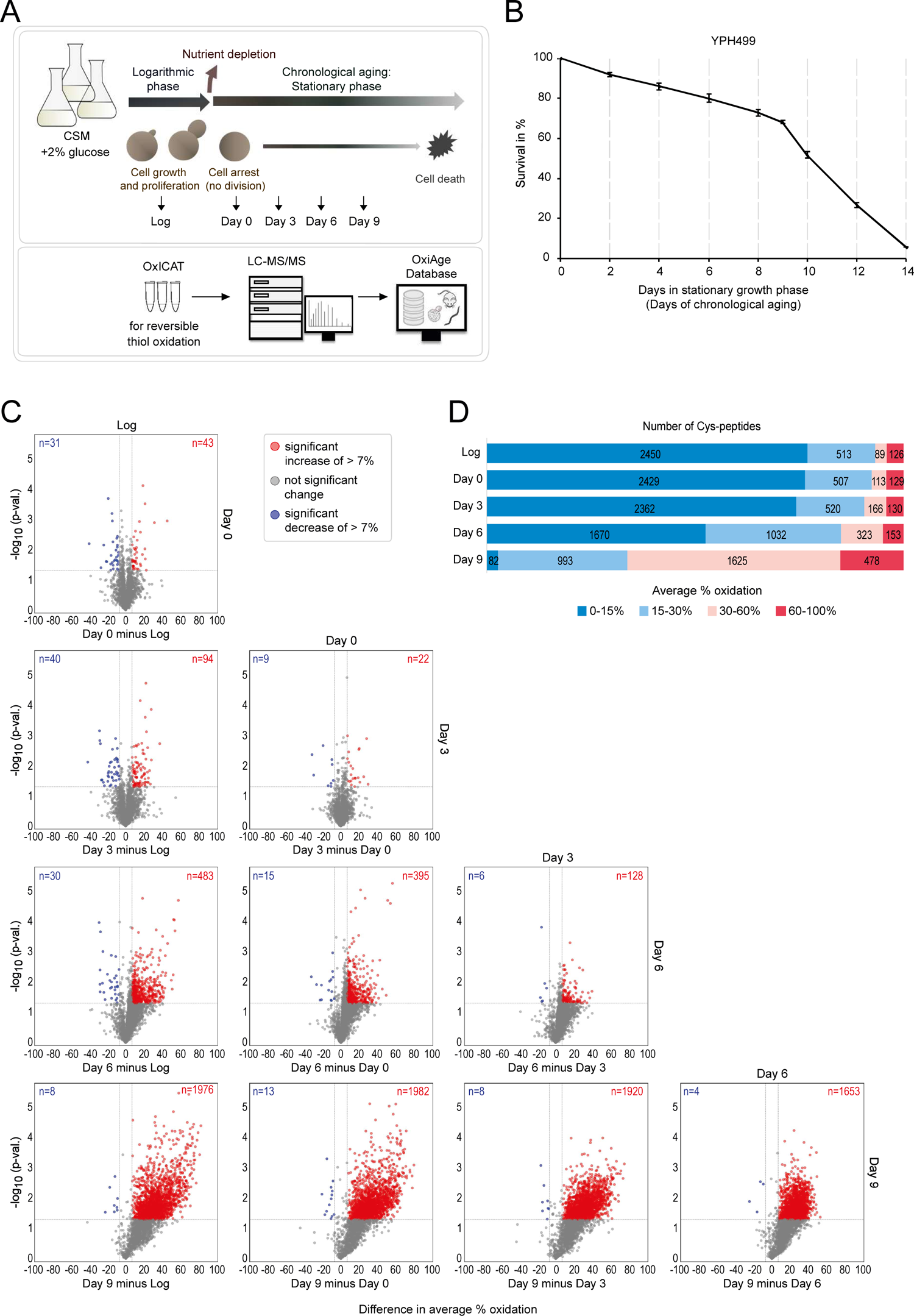
Redox state of yeast proteome changes gradually during progression through chronological ageing. **(A)** Simplified scheme of the experimental design and data analysis. Yeast cells grown in a complete supplement mixture (CSM) medium with 2% glucose were collected during the mid-logarithmic growth (Log) and four consecutive time points upon entry into the stationary phase. Entry into the stationary phase on day 0 marks the start of yeast chronological ageing. Collected samples were subjected to chemical labelling of reversibly oxidized thiols using the OxICAT technique followed by tryptic digestion for LC-MS/MS analysis. Identified peptides were quantified and the proportion of reversibly oxidized cysteine residues (% oxidation) was calculated. The resulting “yeast OxiAge” dataset was compared to data on age-dependent oxidation in other species. A web application was developed containing a comprehensive database of evolutionarily conserved proteome redox modifications during ageing. Upper panel, samples collection; lower panel, samples and data processing. **(B)** Lifespan curve of chronologically aged wild-type yeast *S. cerevisiae*, strain YPH499. Samples were collected on indicated days after entry into the stationary growth phase. Cell viability was determined with propidium iodide (PI) staining and the percentage of dead (stained) cells was determined by FACS. At least two biological replicates per time point were analysed. Data are presented as mean ± SD. **(C)** Pairwise comparison of *in vivo* cysteine modification during proliferation (Log) and different stages of chronological ageing (days 0, 3, 6, and 9) from the “yeast OxiAge” dataset (imputed values). Difference in % oxidation between time points is shown on the x-axis; -log_10_ *p*-value (two-tailed Welch’s *t*-test) is shown on the y-axis.; grey horizontal line, *p*-value = 0.05. Grey dots, peptides of non-significant change in % oxidation or change of less than 7%; blue dots, peptides of a significant decrease in oxidation of more than 7%; red dots, peptides of a significant increase in oxidation of more than 7%. **(D)** Distribution of the *in vivo* oxidation status of 3,178 unique Cys-peptides containing reversible-oxidized cysteine residues from the “yeast OxiAge” dataset (non-imputed values). Peptides from each time point were classified into four oxidation groups based on the average % oxidation. Unique peptides per group were counted for each time point.

To identify peptides with a significant change in oxidation, we compared the levels of thiol oxidation of quantified peptides between each time point of the experiment (Figure 1C). We refer to the proportion of reversibly oxidized cysteine residues as “% oxidation” or “oxidation level”. Minor changes in oxidation level occurred until day 3 of ageing, with a comparable number of peptides exhibiting an increased or decreased % oxidation. On day 3 of ageing compared with the log growth phase, 94 Cys-peptides exhibited a significant increase in oxidation of > +7% (Figure 1C, in red), whereas 40 Cys-peptides showed a significant decrease of > −7% oxidation (Figure 1C, in blue). At day 6 of ageing a visible shift towards higher oxidation of cysteine thiols occurred. At least 16 times more Cys-peptides showed a significant increase rather than a decrease of > 7% in oxidation between day 6 and earlier days of ageing. This shift was even more pronounced on day 9, concomitant with a rapid increase in cell lethality (Figure 1B). The manual classification into the four groups of oxidation levels, 0-15%, 15-30%, 30-60%, and 60-100%, revealed that the basal level of oxidation in the logarithmic phase is below 15% for 2,450 peptides. This confirms previous analyses performed in proliferating yeast demonstrating that in unstressed environmental conditions cysteine residues are present to a large extent in a reduced state (Topf et al., 2018). 126 Cys-peptides had an oxidation of more than 60% in the Log growth phase and this number is comparable with days 0 and 3, with a mild increase at day 6 (Figure 1D). Most changes occur for peptides with an oxidation state above 15% and between 30-60%. We conclude that thiol-containing peptides that are oxidized at a low level in young proliferating cells exhibit a visible shift to a higher redox state during the early days of ageing. Between the young cells and early-aged ones, the shift was observed mostly from low oxidation (< 15%) to medium oxidation (15-60%). Thus, changes in oxidation during ageing were mostly gradual. Only at the late phase of ageing (day 9), the number of highly oxidized peptides increased dramatically, which is likely accompanied by oxidative damage.

### Classification of redox changes in the ageing proteome

To distinguish between different trends in thiol oxidation and identify proteins that are most sensitive to the age-dependent increase in oxidative stress, we performed a clustering analysis of our time-resolved thiol oxidation data. Ten groups (cluster A-J) of thiol-containing peptides are distinguished based on the average oxidation at the different time points (Figure 2, Supplementary Figure 2A and B, Supplementary Table 3). Most of the time points per cluster contain data with an interquartile range IQR < 7, which indicates small variability between grouped peptides per cluster (Supplementary Figure 2B).

**Figure 2.**
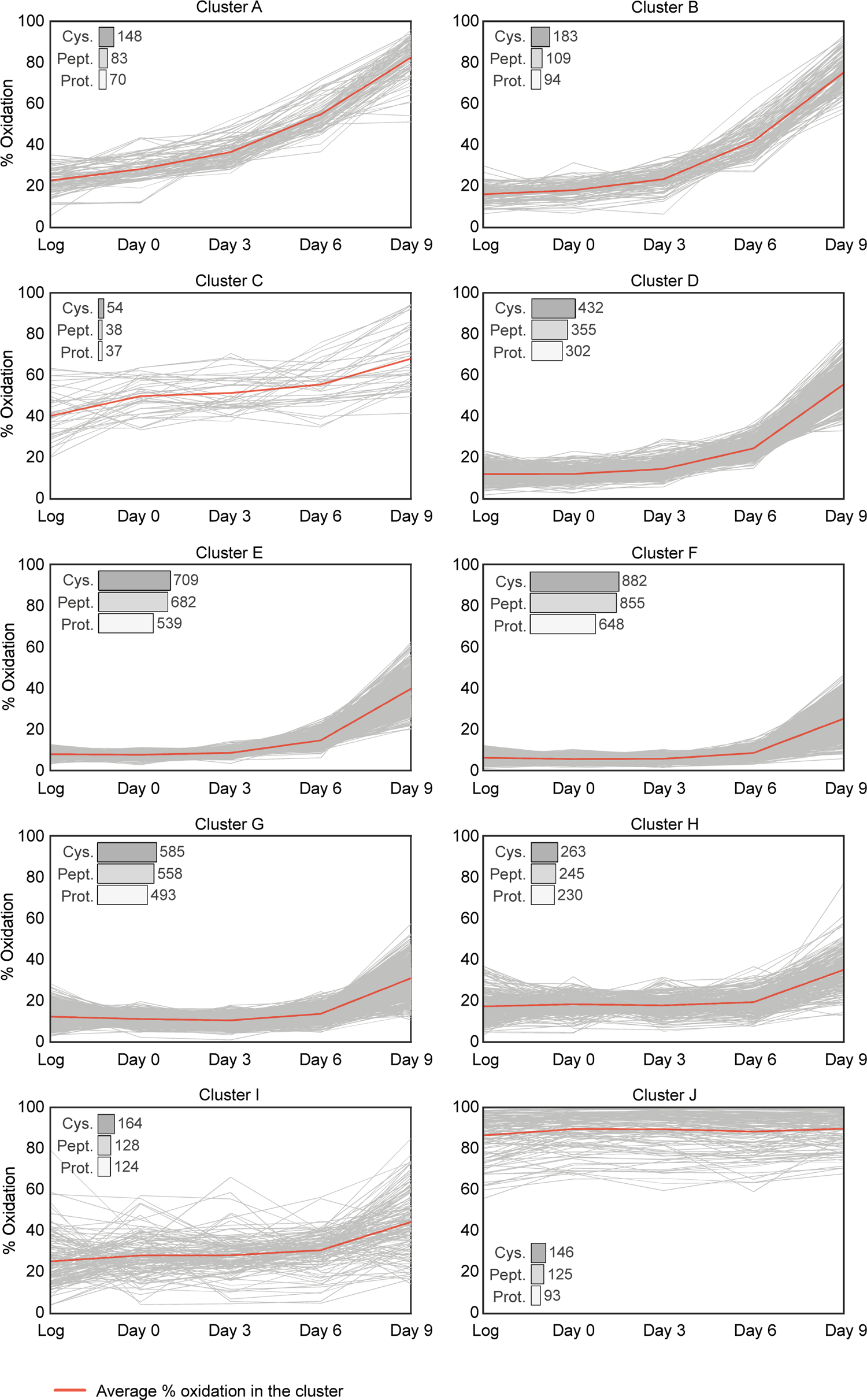
Classification of temporal changes in proteome oxidation based on peptide-specific patterns. Cys-peptides from the “yeast OxiAge” dataset were clustered *in silico* into ten groups exhibiting distinct oxidation patterns. Clusters are named A-J. Peptides within clusters A-C exhibit the fastest increase in oxidation during the early stages of ageing (days 0 and 3). Cluster D contains peptides with average % oxidation increasing during the mid-point of ageing (day 6). Clusters E-H contain peptides with distinct elevation in % oxidation during late ageing (day 9). Peptides from cluster J do not change their high oxidation levels throughout the time course. Cluster I contains peptides with various redox patterns that could not be automatically classified into other groups. Grey traces, average % oxidation for each Cys-peptide and time point; red trace, average % oxidation in the cluster. *Log*, logarithmic phase. *Cys.*, the number of unique cysteine residues within the cluster; *Pept.*, the number of unique Cys-peptides within the cluster; *Prot.*, the number of unique proteins within each cluster. Note that each Cys-peptide can belong only to one cluster; each cysteine can belong to more than one cluster if detected in more than one Cys-peptide; each protein can belong to more than one cluster if the protein has more than one Cys-peptide quantified and grouped within different clusters.

Clusters A, B, C, and D contain peptides with cysteine residues showing a considerable increase in % oxidation at day 3 and day 6 when compared to the Log phase. These clusters contain in total 585 Cys-peptides of 458 unique proteins. The earliest oxidation increase is observed in clusters A, B, and C. Cluster A contains 83 peptides, corresponding to 148 unique cysteine residues and 70 unique proteins (Figure 2). These peptides are significantly more oxidized on day 3 than during the Log phase (corr. *p*-val. < 0.00005, Dunn’s), with an increase from ∼20% to ∼35% and a further increase on day 6 to average oxidation of ∼55% (corr. *p*-val. < 0.0000005, Dunn’s) (Supplementary Figure 2B). For cluster B, Cys-peptides showed a delay in increase of % oxidation compared to cluster A, but globally a significant increase is observed already on the third day of ageing (corr. *p*-val. < 0.00005, Dunn’s). Intriguingly, the earliest increase in % oxidation was seen for peptides in cluster C. This cluster contained Cys-peptides that, on average, were oxidized >40% during the Log growth phase. Although an early increase in oxidation levels is observed, there was a strong variation in % oxidation between Cys-peptides within cluster C (Supplementary Figure 2A).

The majority of thiol-containing peptides group into clusters E-H, with no or little changes in % oxidation during early time points (Supplementary Figure 2A and B). On average, the oxidation of these peptides was < 20% during early and mid-point ageing. Cys-peptides from cluster G exhibited a decrease in oxidation when entering the stationary phase of day 0, and then sequentially increase in the later days. The maximum thiol oxidation in clusters E-H on day 9 was on average < 50%. Thus, protein thiols within these clusters can be considered less sensitive to oxidation-dependent modifications during ageing.

Several peptides grouped into a non-homogeneous cluster I, containing 128 Cys-peptides in 124 proteins, 55 of which were unique to this cluster (Figure 2). In general, cluster I consisted of Cys-peptides that could not be grouped into any other cluster and did not exhibit any consistent trend (Supplementary Figure 2A).

Custer J contains 125 Cys-peptides corresponding to 93 proteins that did not change their redox status and remained highly oxidized (> 85%) over the time course (Figure 2). 80 proteins contained at least one Cys-peptide assigned exclusively to cluster J, while 13 proteins contained peptides allocated additionally in clusters C-I. Intriguingly, no protein with Cys-peptides assigned to clusters A or B was present in cluster J.

In summary, whereas most protein thiols within the “yeast OxiAge” dataset were oxidized late during ageing, we identified proteins that contained cysteine residues sensitive to changes in oxidation during the early phase of ageing. We hypothesize that these proteins are likely to influence cellular response and organismal fate during biological ageing.

### Cysteine residues oxidized during early ageing exhibit a common amino acid signature

We asked whether the early-oxidized Cys-peptides exhibit any significant similarities that might affect their high sensitivity to oxidation. We analysed the local amino acid environment of the early-oxidized cysteine residues (Supplementary Table 3). We found that a common feature of the peptides assigned to clusters A, B, and C was the presence of additional cysteine residue surrounding the early oxidized one (Figure 3A). Within Cys-peptides that showed an increase in oxidation on days 0 and 3 of ageing an additional cysteine in close proximity to the oxidation-sensitive cysteine residue was overrepresented in the peptide sequence. A second cysteine was present either at position −3 or +3 from the oxidized cysteine subjected to motif analysis. We found that 44 out of 83 Cys-peptides in cluster A contained two oxidized cysteine residues that both formed the CXXC motif. Subjecting both cysteines to the motif analysis led to the generation of a symmetric signature, where the same oxidized cysteine might be present either at position 0, −3 or +3, depending on the analysed residue (position 0). In case of 17 Cys-peptides, a single cysteine from the CXXC motif was found to be early oxidized within cluster A (among them for two proteins both cysteine residues were found within two different Cys-peptides). Interestingly, two proteins, the stress response protein, Nst1, and the aminopeptidase, Fra1, showed a CXXCXXC motif with all three cysteine residues present within the same early oxidized Cys-peptide. The CXXC motif was also found for Cys-peptides in clusters B and C, but a cysteine residue present at either the position −3 or +3 occurred with a lower frequency than in cluster A (Figure 3B). The CXXC motif has been previously identified for cysteine residues responding to the treatment with H_2_O_2_ with a significant increase in oxidation (Topf et al., 2018) and during ageing (Brandes et al., 2013). Topf *et al*. noted that this motif was also associated with the binding of zinc ions. Our analysis confirmed the significant overrepresentation of the CXXC motif in zinc-binding proteins, similar to early-oxidized proteins from clusters A-C (Figures 3C and D).

**Figure 3.**
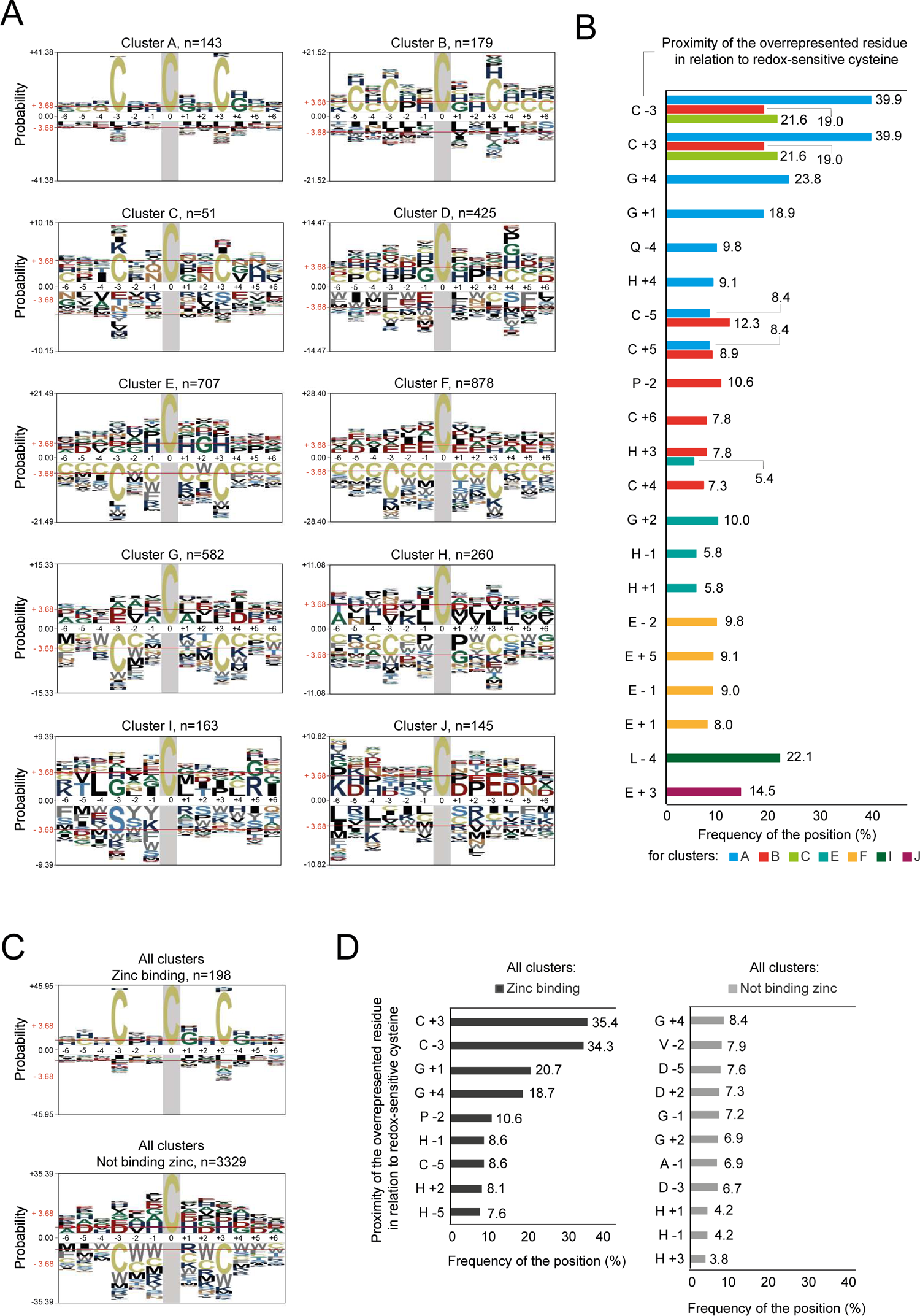
Motif analysis of oxidized cysteine sites demonstrates a conserved sequence in individual clusters. **(A)** Consensus motifs for modified cysteine residues for peptides within each cluster A-J. Clusters A, B and C show a significant overrepresentation of cysteine residues at either position −3 or +3, while clusters E-H shows significant selection against cysteines at the same positions. Graphs present the log-odds binominal probabilities. The sum of the log-odds probabilities for the largest column is used as the maximum value for the y-axis. Residues are scaled proportionally to their log-odds binominal probabilities under the chosen background with the biggest residue indicating the strongest statistical significance. Red horizontal line at +/-3.68, *p*-value = 0.05 (Bonferroni corrected). Probabilities above +3.68, overrepresented statistically significant residues; probabilities below −3.68, underrepresented statistically significant residues. Amino acid residues are coloured according to their physicochemical properties (see the **Methods** section). Grey area, fixed position of the oxidized cysteine residue. Column numbers, positions from the centred oxidized cysteine at position 0. *n*, number of unique motifs analysed in individual clusters. **(B)** Frequency in % of the significantly overrepresented amino acid residues at specified positions in relation to the oxidized cysteine within the individual clusters. Only clusters A, B, C, E, F, I and J contain significantly enriched motifs. Columns indicate the residue (*C*, cysteine; *G*, glycine; *Q*, glutamine; *H*, histidine; *P*, proline; *E*, glutamic acid; *L*, leucine) and its position relative to the oxidized cysteine. **(C)** Consensus motifs for zinc-binding and not-binding redox-sensitive cysteines within all clusters. Significant overrepresentation of the CXXC motif is observed for zinc-binding residues. **(D)** Frequency in % of the significantly overrepresented amino acid residues at specified positions in relation to the oxidized zinc-binding or not-binding cysteine. Columns indicate the residue (*C*, cysteine; *G*, glycine; *H*, histidine; *P*, proline; *D*, aspartic acid; *A*, alanine; *V*, valine) and its position relative to the oxidized cysteine.

Early oxidized Cys-peptides in cluster A have a tendency for the enrichment of a simple non-polar amino acid glycine (Gly/G) at positions +1 and +4 with a frequency of ∼20% (Figure 3B). Smaller frequency is observed for a positively charged histidine at position +4 (His/H; frequency ∼9%) and polar glutamine at position −4 (Glu/Q; frequency ∼10%). Cysteine residues in cluster B show a tendency for more complex motifs composed of several additional Cys in close proximity: 3, 4, 5 or 6 positions away from the oxidized cysteine (Figure 3A; Cluster B). Peptides grouped in cluster D, which showed mid-point oxidation during ageing, did not exhibit a specific motif (Figure 3A; Cluster D). Interestingly, Cys-peptides within the clusters E-H, which were oxidized during late ageing, showed a significant underrepresentation of motifs with multiple cysteine residues in close proximity. Although overrepresentation of any particular amino acid sequence was less common in these clusters, cluster F contained a significant enrichment for glutamic acid in close proximity to late-oxidized cysteine residues (Glu/E; position ±1; frequency ∼20%) (Figure 3B). Here, an additional acidic residue at positions −2, and −5 (frequency < 10%) was found. Interestingly, the trend of selection for acidic amino acid was also observed in the “yeast OxiAge” dataset for peptides of high redox modification independent of cellular age (cluster J), with enrichment for glutamic amino acid and close to the significant overrepresentation of aspartic acid (Asp/D) (Figures 3A and B; Cluster J).

The identification of unique cysteine motifs of peptides that exhibit higher sensitivity to oxidative modification during early ageing might serve as a starting point for the identification of potential residues that respond to oxidative stress related to ageing.

### Peptide classification reveals distinct functional groups of early-oxidized proteins

To identify the function of the proteins within the distinct clusters, we analysed their annotated subcellular localization (Figure 4A, Supplementary Table 3) and performed a Gene Ontology (GO) enrichment analysis for biological processes (BP) and molecular functions (MF) (Figure 4B and C, Supplementary Figure 3A and B, Supplementary Table 4). We did not observe any correspondence between peptide clusters and subcellular compartments. An exception was cluster J enriched in proteins localised in the endoplasmic reticulum (ER), Golgi apparatus, mitochondrion, and vacuole (Figure 4A). Peptides within clusters C and J were characterized by exceptionally high oxidation states already in proliferating cells. Indeed, the cellular organelles mentioned above have a higher redox potential compared with, for example, the cytoplasm (Go & Jones, 2008). A large part of early oxidized Cys-peptides belonging to cluster A corresponded to proteins allocated to ribosomes. This is consistent with GO enrichment analysis showing a significant enrichment in cytoplasmic translation processes (Figure 4B) and translation activities (Figure 4C) within this cluster. An example of an early oxidized protein regulating cytoplasmic translation is Tif35 (human eIF3g), a subunit of translation initiation factor eIF3. One quantified peptide contained two cysteine residues at positions 112 and 121, which form a disulfide bond. Change in % oxidation of this Cys-peptide was fast with an increase from 25.8 ± 8.7% on day 0 to 42.4 ± 5.4% on day 3. tRNA synthetases were also found within cluster A. Among them was Mes1 (human MetRS/MARS1) with two cysteine sites oxidized during the early stages of ageing. Cys353 (cluster A) had a basal oxidation level of 22 ± 4.5% during the Log phase that elevated to 37.8 ± 5% on day 3. The consecutive time points followed this trend with an increase to 62 ± 11.7% and 92.9 ± 4.8% on days 6 and 9, respectively. As suggested by structure prediction [AF-P00958-F1, AlphaFold prediction (Jumper et al., 2021)], this residue is localised on the surface of the protein and interacts with three other residues at positions 337, 340, and 350. The second thiol was found in Cys321 and was oxidized slower than the other cysteine (cluster D).

**Figure 4.**
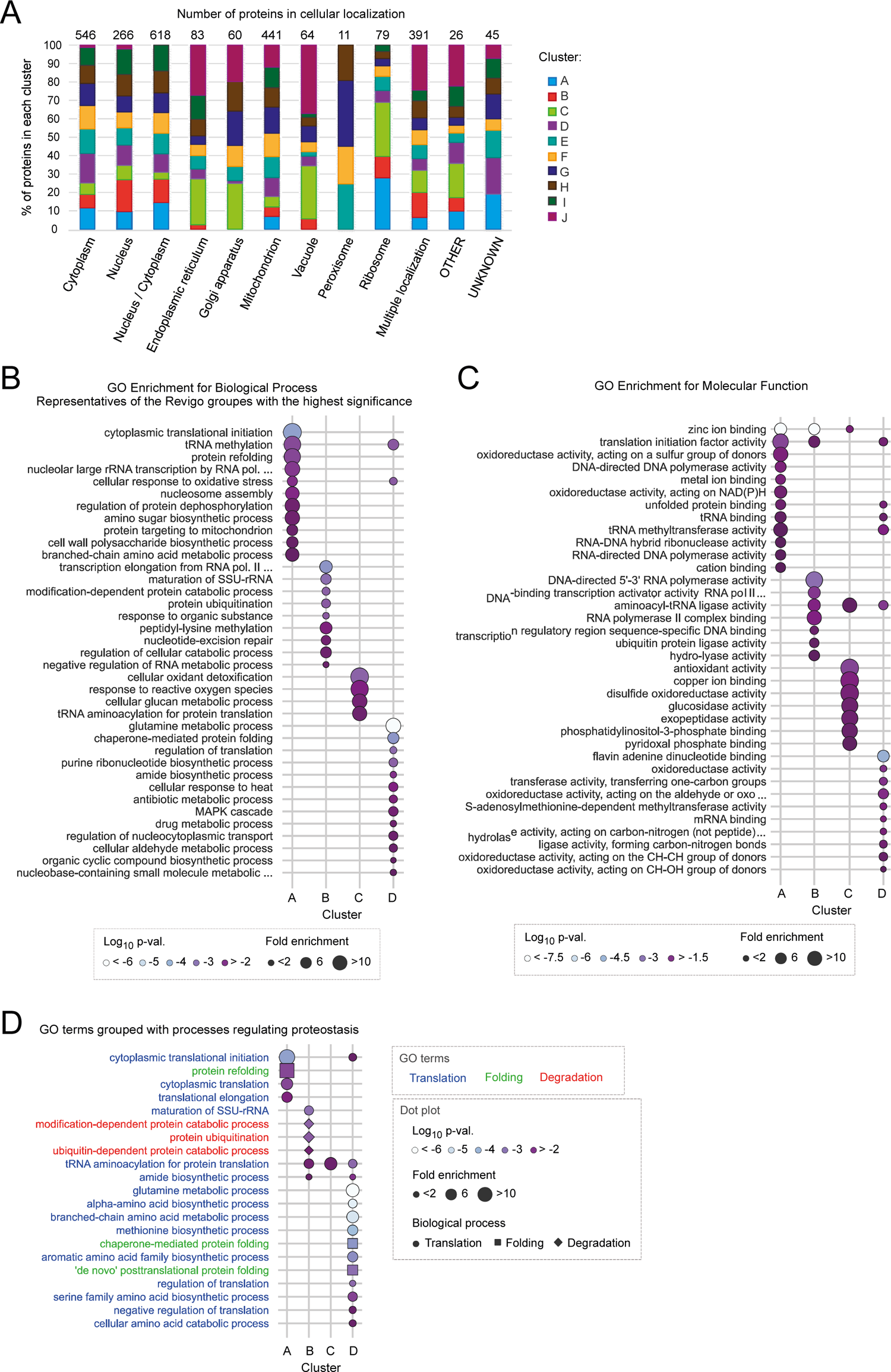
Distinct functional groups of proteins oxidized at different stages of ageing. **(A)** Distribution of proteins with at least one Cys-peptide belonging to individual clusters A-J according to subcellular localization. The frequency of proteins within a particular cluster is presented for each localization. Gene ontology (GO) terms for cellular components were retrieved from UniProtKB (see the **Methods** section). Total number of proteins per localization is indicated above the bar plots. **(B)** GO term enrichment analysis for biological process (BP) for individual clusters A-D. For simplification of the graph GO BP terms were grouped by Revigo based on the relation between GO terms. GO BP terms shown are representative of the Revigo groups with the highest statistical significance (Fisher’s test, Benjamini-Hochberg correction). A maximum of 15 best-scored terms are shown. Dot size, fold enrichment; dot colour, log_10_ uncorrected *p*-value. **(C)** GO term enrichment analysis for molecular function (MF) for individual clusters A-D. A maximum of 15 best-scored terms based on the highest statistical significance (Fisher’s test, Benjamini-Hochberg correction) are shown. Dot size, fold enrichment; dot colour, log_10_ uncorrected *p*-value. **(D)** All GO BP terms for clusters A-D that represent processes involved in proteostasis regulation or are grouped in the same Revigo group as the terms representing the proteostasis regulation (see the **Methods** section). Colours of GO term indicate the proteostasis process: blue, cytoplasmic translation; green, protein folding; red, protein degradation. Marker size, fold enrichment; marker colour, log_10_ uncorrected *p*-value; marker shape, biological process of proteostasis: circle, cytoplasmic translation; square, protein folding; diamond, protein degradation.

Further, we found that the biological processes “protein refolding” and “ribosomal RNA (rRNA) transcription by RNA polymerase I” were strongly enriched in cluster A (Figure 4B). The “yeast OxiAge” dataset identified six unique cysteine sites of RNA polymerase I subunit Rpa135 (human RPS135/POLR1B). Cys1104 and Cys1107 were found in two different peptides grouped into cluster A, with oxidation of ∼20% in proliferating cells, later increasing to ∼30% on days 0 and 3 and > 44% on day 6. Cysteine residues at positions 158, 584, 585, and 734 were grouped into cluster D with an increase in oxidation on days 3 and 6. Another protein with early oxidized cysteine residues (positions 40, 43, 68 and 73) is Mig1, involved in glucose repression. Activation of Mig1-repressed genes is critical for maintaining chronological ageing in yeast (Maqani et al., 2018). It is strictly regulated by the phosphorylation of multiple serine residues, but the possibility of its regulation through cysteine residue modification has so far been undescribed.

Contrary to cluster A, cluster B contains proteins mostly found to regulate protein degradation and mRNA transcription (Figure 4B). An example of a protein involved in the latter process is a subunit of RNA polymerase II, Rpb9 (human RPB9/POLR2I) with a single thiol group on Cys32 oxidized from 16.6 ± 4.1% during Log to 34 ± 7.1% on day 3. This fast oxidation was followed by a continuous increase to 44.8 ± 3.9% and 71.9 ± 12.6% on consecutive days. Consistently, a lower increase in % oxidation until the mid-point of ageing (< 60%) might indicate a possible oxidation-dependent regulation of the protein during ageing.

Cluster C was enriched in proteins involved in response to oxidation, glucan metabolism, and tRNA aminoacylation for translation (Figure 4B). A prominent example is the glutathione-dependent disulfide oxidoreductase Grx1 (GLRX and GLRX2 in humans), which activity is involved in protection against oxidative damage (Luikenhuis et al., 1998). Cys27 and Cys30 are present in a single peptide with strong oxidation of more than 40% in proliferating cells, and subsequent increase on day 0 to ∼50% with no significant change on day 3. A strong increase to more than 70% was observed on day 6. Modification of both sites was previously reported (Hakansson & Winther, 2007; Yu et al., 2008). Cys27 is located in the active site of the oxidoreductase, suggesting a direct effect on Grx1 activity. Mutation of Cys30 to serine was experimentally shown to increase the activity of this oxidoreductase (Discola et al., 2009).

Cluster D was significantly enriched with proteins involved in glutamine metabolism, but also in chaperone-mediated protein folding and regulation of translation (Figure 3B). Similar to cluster A, proteins linked to translation initiation factor activity, tRNA binding, and unfolded protein binding were enriched in cluster D (Figure 3C).

Proteins with peptides identified in clusters E-H were enriched for similar biological processes (Supplementary Figures 3A and B). For example, proteins in clusters E and F were significantly enriched for a global response to oxidative stress, tRNA aminoacylation, amino-acid biosynthesis, and ribosomal biogenesis. This included some ribosomal proteins, such as Rps11B (Cys58 and Cys128) or Rps8B (Cys179; Cys168 in cluster G). We identified also a variety of regulators of protein folding, such as chaperonin-containing TCP-1 (CCT) multiprotein complex components or heat shock proteins (HSP). This characteristic was shared between clusters E, F, and G. Additionally, mitochondrion organization and metabolic processes were among the significantly enriched biological processes.

Cys-peptides in cluster I and J correspond to proteins involved in a variety of biological processes relevant to cellular metabolism (Supplementary Figure 3A). The analysis of the cellular compartmentalization of proteins grouped in cluster J revealed that they are preferentially localised in the mitochondrion (16 proteins) and vacuole (13 proteins) (Figure 4A). Accordingly, proteins assigned to cluster J are involved in protein targeting to the mitochondrion, cell wall organization, glycosylation, and general regulation of intercellular transport (Supplementary Figure 3A). An example of such a protein is the vacuolar aspartyl protease Pep4. We found three cysteine residues at positions 122, 127, and 328, oxidized above 80% over the yeast’s lifespan. All oxidized cysteine sites are localised within the peptidase A1 domain and are involved in disulfide bond formation. Another example is the mitochondrial intermembrane space import and assembly protein 40 (Mia40) (Chacinska et al., 2004). For this protein, Cys317 identified in this study is one of six known cysteine sites involved in the formation of disulfide bonds.

In agreement with the motif analysis, we found a significant enrichment for zinc-binding proteins in clusters A, B, and C (Figure 3C). We did not observe this enrichment in any other cluster of Cys-peptides oxidized at later stages of ageing (Supplementary Figure 3B). Notably, clusters C and J containing strongly oxidized peptides already during the proliferation stage were significantly enriched in proteins known for binding copper ions (Supplementary Figure 3B).

The functional analysis of proteins carrying redox-sensitive thiols revealed a remarkable distinct pattern. It appears that proteins involved in key cellular processes such as translation, folding and degradation contain cysteine residues that are preferably oxidized during the early stages of ageing.

### Oxidation of proteins regulating proteostasis as an early response to ageing

Protein homeostasis (proteostasis) depends on the coordinated interplay of protein production, folding, and degradation. Loss of protein homeostasis is a longstanding determinant of the ageing process (Hohn et al., 2017; Kaushik & Cuervo, 2015). Analysis of GO terms associated with proteostasis demonstrated the strongest overrepresentation of these processes in cluster A of early oxidized proteins, with the highest fold enrichment for cytoplasmic translational initiation (fold enrichment > 12; corr. *p*-val. = 0.009, Fisher’s) and protein refolding (fold enrichment > 8; corr. *p*-val. < 0.03, Fisher’s) (Figure 4D). Moreover, early oxidized proteins in cluster B were found to be significantly enriched in protein ubiquitination and degradation terms (fold enrichment > 2.5; corr. *p*-val. < 0.05, Fisher’s). Since our functional analysis revealed that age-dependent oxidation might play a role in all three processes, we wanted to further determine the identity of the associated redox-sensitive proteins. We performed a search of GO terms and assigned all proteins to three groups: cytosolic translation, protein degradation, and protein folding (see the Methods section). In total, ∼16% of all proteins were annotated as involved in proteostasis (Figure 5A, Supplementary Table 5). Cys-peptides of these proteins exhibited a similar oxidative shift towards higher values as cells progressed through ageing compared with peptides belonging to other processes. Notably, a strong preference for increased oxidation during early ageing was observed (Supplementary Figure 4). Although the total number of proteostasis-regulating proteins was higher in bigger clusters (E-H), a higher percentage of proteins was associated with the smaller clusters A-C (Figure 5A). Thus, we focused our analysis on proteins within cluster A-D which contained proteins with thiols oxidized early during ageing. Cys-peptides within the clusters were modified at a similar level during the analysed time points. The oxidation levels of Cys-peptides were independent of the involvement of the protein in a particular proteostasis-associated process (Figure 5B). In cluster A, the majority of proteostasis-regulating proteins were functionally related (Figure 5C). Here, we distinguished two main groups: i) cytosolic ribosomal proteins and ribosome-associated proteins, and ii) proteins involved in protein folding. Most striking is the enrichment in core ribosomal proteins including Rpl37A/B, Rpl43B, and Rps29A/B, as well as other translation-regulating proteins, such as Gis2, Yef3, Tif5 and Sui3. All these proteins were previously identified as redox-sensitive in response to exogenous oxidative stress in different species (Brandes et al., 2011; Menger et al., 2015; Su et al., 2019; Topf et al., 2018). In addition, we identified subunits of translation initiation factors, such as Tif35 and Gcd11, and proteins involved in degradation, namely Rqt4, Hrt1 and Hel2. Rqt4 and Hel2 are both part of the ribosome-associated quality control triggering complex (RQT). The complex acts during ribosome stalling and contributes to the dissociation of the ribosome (Matsuo et al., 2017; Matsuo et al., 2020). Further, four co-chaperones were identified that are necessary for the import of precursor proteins into the mitochondria, namely Ydj1, Xdj1, Mdj1, and Zim17. While Mdj1 and Zim17 are localised in the mitochondrial matrix (Burri et al., 2004; D’Silva et al., 2003), Ydj1 and Xdj1 are cytosolic proteins that were found to deliver mitochondrial precursor proteins to the organelle (Opalinski et al., 2018; Xie et al., 2017).

**Figure 5.**
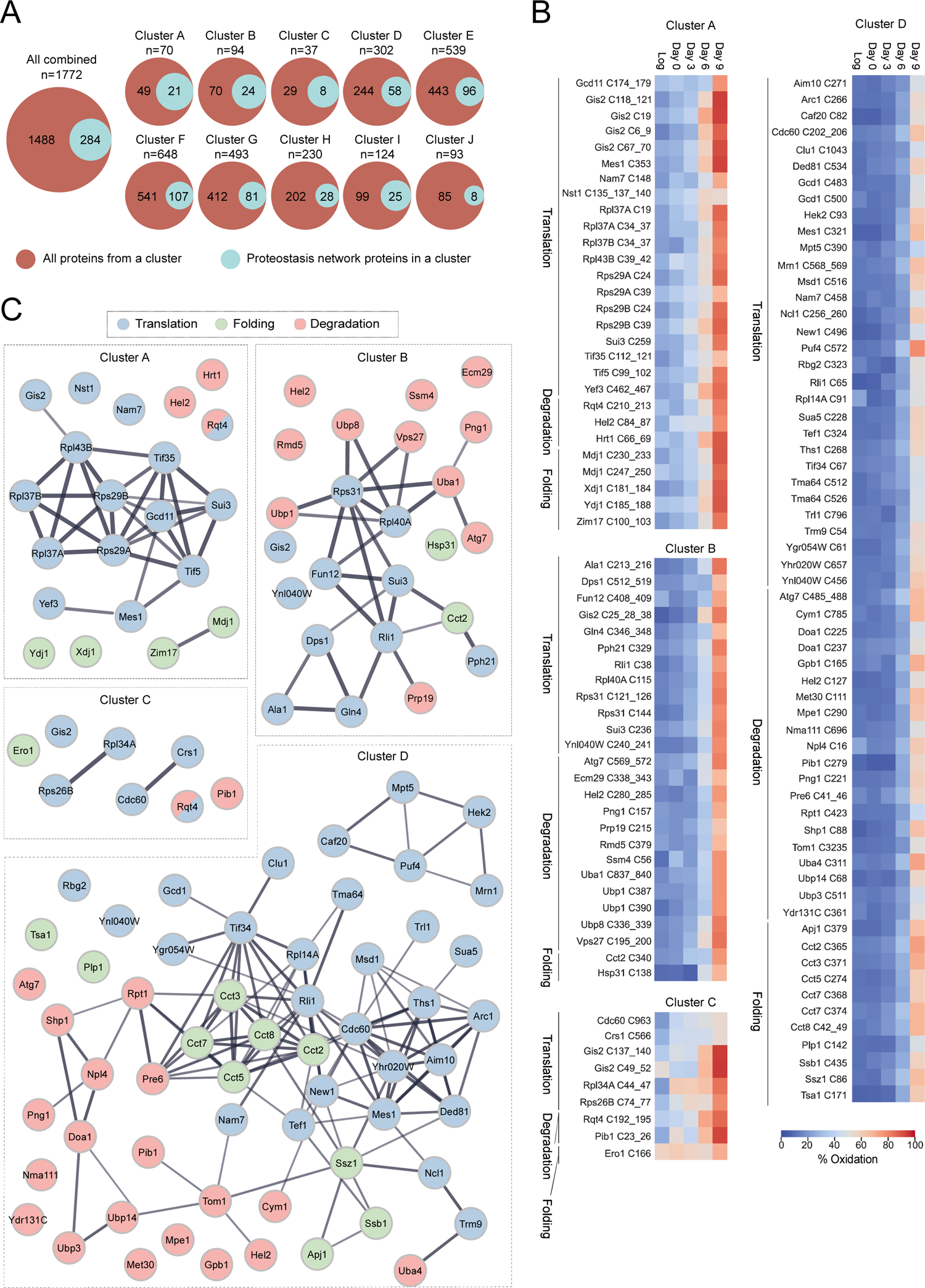
Proteostasis-regulating proteins exhibit progressive oxidation patterns. **(A)** Venn diagram of the number of proteins involved in proteostasis (turquoise) among all proteins that have at least one Cys-peptide grouped within individual clusters (red). *n*, total number of all proteins within the cluster. **(B)** Heatmaps present the average % oxidation of Cys-peptides of proteostasis-regulating proteins from early- and mid-oxidized clusters A-D. Each Cys-peptide is presented in the form of cysteine ID (CysID): name of the protein followed by residue name (cysteine, *C*) and position/s of the quantified residue/s within the peptide. Peptides are grouped in alphabetical order and by the biological process related to proteostasis (translation, degradation, folding). For proteins that belong to more than one process, a representative one was selected (see **Supplementary Table 3**). **(C)** STRING protein association network for proteostasis-regulating proteins for individual clusters A-D. Nodes, proteins; lines, interaction evidence. Line thickness, the strength of the evidence. Nodes are coloured based on the biological process the protein is involved in. Proteins involved in more than one process have more than one colour.

Consistent with the GO enrichment analysis (Figure 3A), proteins with cysteine residues grouped within cluster B are mostly associated with proteome degradation. Thus, the centre of the network is formed by ubiquitin-containing ribosomal proteins Rps31 and Rpl40A with branches extending to proteins involved in translation and degradation (Figure 5C). About half of the proteins of proteostasis-regulating proteins within cluster B are associated with protein degradation. This includes the ubiquitin-specific proteases Ubp1 and Ubp8, the ubiquitin-activating enzyme Uba1 and E3 ubiquitin ligases Hel2 and Ssm4. Many enzymes involved in the ubiquitin-proteasome system are sensitive to oxidative stress and oxidation of cysteine residues in the catalytic domains inactivates the enzymes (Shang & Taylor, 2011). Importantly, the redox-regulated activity of deubiquitinating enzymes of the cysteine protease family contributes to the balance of ubiquitinated protein species in the cell.

Besides proteins involved in the ubiquitin-proteasome system (UPS), several proteins with peptides found in cluster B are involved in cytoplasmic translation. However, in contrast to cluster A, only a few are located within the ribosome itself (Supplementary Table 3; Peptides and Localization). Among the proteins involved in the translation process are the translation initiation factor Sui3, tRNA synthetases (Dps1, Ala1, Gln4) and proteins involved in ribosome biogenesis (Fun12, Rli1). It is largely unknown how these factors are regulated by oxidative thiol modification. It was shown that oxidation of tRNA synthetases impairs editing activity and consequently decreased translation fidelity due to the incorporation of wrong amino acids during polypeptide chain production (Ling & Soll, 2010).

Cluster C contains 37 proteins, out of which 6 are involved in cytosolic translation. Among them are two ribosomal proteins, Rps26B and Rpl34A, and two tRNA synthetases, Cdc60 and Crs1/YNL247W (Figure 5C). Both ribosomal proteins were found to have a single peptide oxidized on average between 20% and 30% with an increase up to even 60% on day 0, stabilizing at this level until day 6 (Figure 5B). Both peptides were also found to be significantly more oxidized in proliferating cells in response to exogenous and endogenous oxidative stress (Topf et al., 2018).

Cluster D contained a total of 29 proteins involved in translation, with only one ribosomal protein, Rpl14A. Other translation-associated proteins formed subclusters including tRNA synthetases (e.g. Cdc60, Msd1, Ths1, Mes1, Ded81), regulators of tRNAs function (e. g. Sua5, Ncl1, Trm9, Arc1), mRNA binding proteins (e.g. Caf20, Puf4, Mrn1, Hek2), and translation initiation factors (e.g. Tif34, Clu1) (Figure 5C). The second most noticeable group of proteins was related to protein folding, specifically *de novo* folding downstream of translation. This included the chaperonin complex TRiC/CCT. This complex consists of eight subunits. We identified five of them within cluster D: Cct2, Cct3, Cct5, Cct7, and Cct8. Moreover, the ribosome-associated chaperones Ssz1 and Ssb1 were quantified. This might indicate that the protein folding machinery is a target of redox regulation during the mid-point of ageing.

Within cluster D, 19 proteins were involved in degradation and they formed a distinct but dispersed group (Figure 5C). The larger subgroup belongs to enzymes of the UPS involved in ubiquitin ligation (E1; e.g. Tom1, Hel2, Pib1), ubiquitin removal (DUBs; Ubp14, Ubp3), and ubiquitin chain binding receptor (Npl4, Shp1). Moreover, the proteasome subunits Rpt1 (ATPase of the regulatory particle) and Pre6 (alpha subunit of the core) were identified. However, the vast majority of proteasome subunits had Cys-peptides grouped in clusters F and G (Supplementary Figure 5) suggesting that the proteasome is a target of oxidation only at the late stages of ageing. This indicates a possibility of either a late regulatory role of the oxidation of proteasome or oxidation-dependent damage.

Our analysis revealed proteins with specific oxidation-sensitive thiols that are essential players during proteostasis. The molecular consequence of the rapid increase in oxidative modification on the specific cysteine residue is for most of the proteins unknown and further research will be necessary to elucidate the consequences for the protein itself and its role during the ageing process.

### Early oxidation of proteostasis-regulating proteins is conserved among species

The increase in reactive oxygen species accompanied by increasing oxidative damage of proteins during ageing is a universal phenomenon also found in higher eukaryotes (Agarwal & Sohal, 1996; Gladyshev et al., 2021; Hohn et al., 2013; Stadtman, 2001). Reversible oxidative modification on the proteome-wide scale during organismal ageing was addressed to the best of our knowledge only in four other datasets: budding yeast *S. cerevisiae* wild-type strain DBY746 (Knoefler et al., 2012), nematode *C. elegans* wild-type strain N2 (Brandes et al., 2013), fruit fly *D. melanogaster* wild-type strain *white Dahomey* (Menger et al., 2015), and in mouse *M. musculus* wild-type strain C57BL/6 (Xiao et al., 2020). We aimed to compare the published datasets (Supplementary Table 6) together with the “yeast OxiAge” compendium for commonly identified proteins with reactive thiol groups (Supplementary Table 2). The analysis was limited by the different methodologies used and the total number of proteins identified in the different studies. Thus, the overlap between all the datasets was rather small (Figure 6A and Supplementary Table 7). We reasoned that most likely early oxidized proteins would have the largest impact on cells’ functions during ageing. Since the decline in proteostasis is a universal hallmark of ageing (Lopez-Otin et al., 2023), we specifically searched in the published datasets (Supplementary Table 6) for the equivalent proteins and cysteine residues that we identified in the “yeast OxiAge” sub-dataset for proteostasis (Supplementary Tables 8 and 9). We speculated that in other organisms similar functional classes of proteins might coordinate the ageing process. Therefore, we pairwise matched only the proteostasis-regulating proteins quantified in the “yeast OxiAge” dataset with the four existing redoxome datasets of aged eukaryotes.

**Figure 6.**
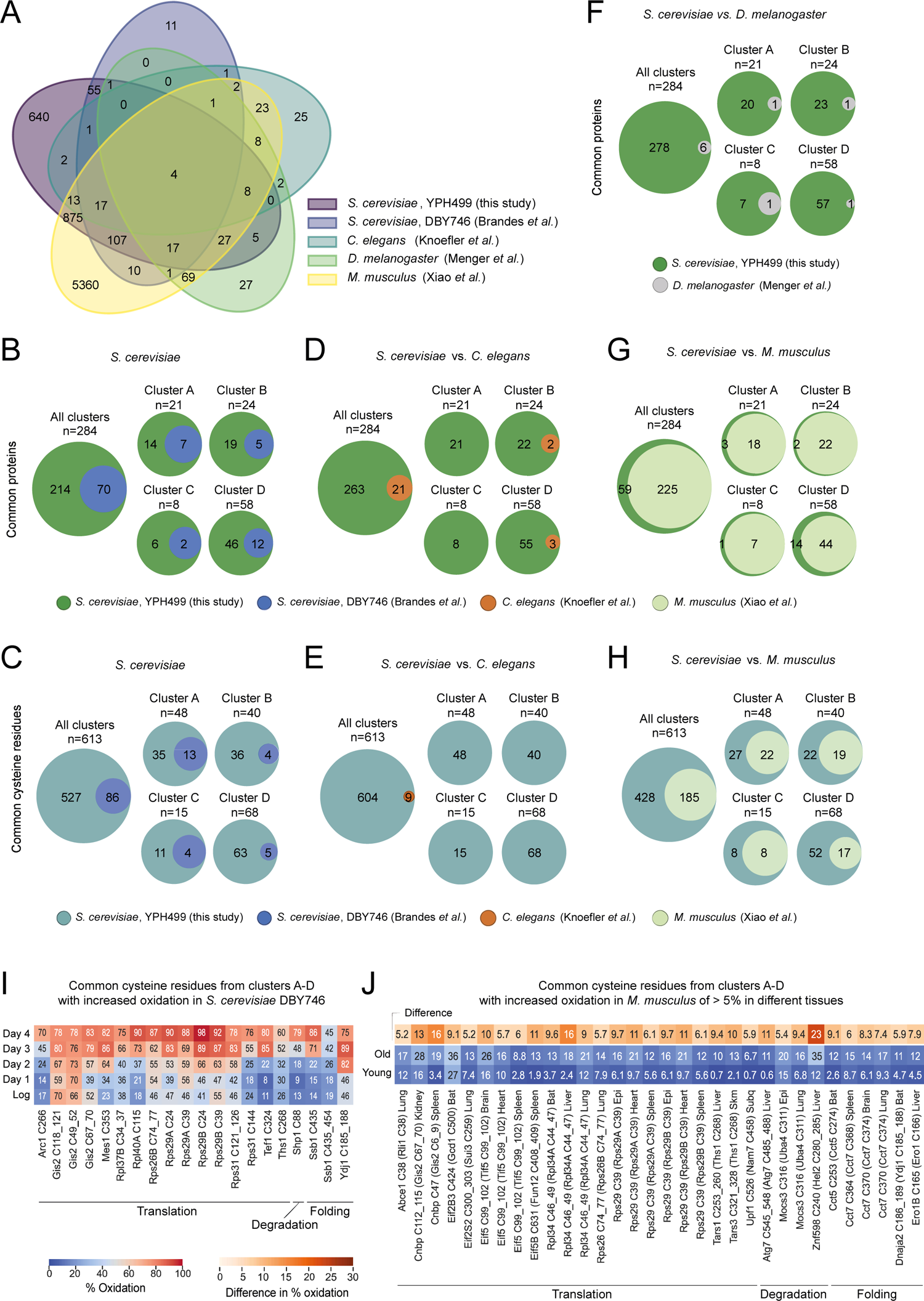
Evolutionarily conservation of oxidation of proteostasis-regulating proteins during ageing. **(A)** Venn diagram of common proteins found oxidized during ageing in different species. Purple, *S. cerevisiae* strain YPH499 (“yeast OxiAge”, this study); blue, *S. cerevisiae* strain DBY746 (Brandes *et al*.); turquoise, *C. elegans* strain N2 (Knoefler *et al*.); green, *D. melanogaster* strain white Dahomey (Menger *et al*.); yellow, *M. musculus* strain C57BL/6 (“Oximouse”, Xiao *et al*.). **(B-H)** Pairwise comparison of commonly oxidized proteins (B, D, F, G) and cysteine residues (C, E, H) between species. Orthologs were compared to the whole “yeast OxiAge” dataset (“All clusters”) and each cluster A-D individually. Green, unique proteins from “yeast OxiAge” dataset (B, D, F, G); turquoise, unique cysteine residues from “yeast OxiAge” dataset (C, E, H); blue, unique proteins or cysteine residues from yeast Brandes *et al*. dataset; brown, unique proteins or cysteine residues from worm Knoefler *et al*. dataset; grey, unique proteins from fruit fly Menger *et al*. dataset; pale green, unique proteins or cysteine residues from mice Xiao *et al*. dataset. **(I)** Heatmap of average % oxidation of Cys-peptides from yeast Brandes *et al*. dataset for commonly oxidized cysteine residues with the “yeast OxiAge” dataset. Only peptides from clusters A-D of “yeast OxiAge” and peptides with increased oxidation in Brandes *et al*. are shown. Peptides are presented in alphabetical order and sorted based on the biological process. **(J)** Heatmap of average % oxidation of Cys-peptides from the “Oximouse” dataset for commonly oxidized cysteine residues with the “yeast OxiAge” dataset. Only peptides from clusters A-D of “yeast OxiAge” and peptides with increased oxidation of > 5% in “Oximouse” are shown. Peptides are presented in alphabetical order and sorted based on the biological process. The difference in average % oxidation between old and young mice is shown in a single-colour warm palette heatmap.

First, we compared the two yeast strains: YPH499 (this study, “yeast OxiAge”) and DBY746 (Brandes et al., 2013). We identified 70 proteins involved in proteostasis as redox-sensitive (Figure 6B) with 86 common cysteine residues (Figure 6C). As expected, the highest number of matching proteins and cysteines was present in the biggest clusters of the late-oxidation, clusters E and F (Supplementary Figures 6A and B, Supplementary Tables 8 and 9). For clusters A-D with proteins showing early- and mid-point increase in oxidation, 26 proteins were identified in both yeast strains (Figure 6B). Among them were ribosomal proteins, such as Rpl37B, Rps26B, or Rps31, regulators of translation, like Yef3, Tef1, and Gis2, as well as tRNA synthetases, i.e. Mes1, Dps1, and proteins involved in folding, i.e. CCT components and Ydj1 (Figure 6I, Supplementary Table 8). Furthermore, 36 specific cysteine residues were quantified in both yeast datasets for proteostasis-regulating proteins from clusters A-D (Figure 6C). Three out of 26 proteins did not have any matching quantified cysteine residue, and this includes Cct8 (“yeast OxiAge” Cys sites 42, 49, 252, 298 and Brandes *et. al*. Cys336), Gcd1 (“yeast OxiAge” Cys sites 356, 370, 483, 500 and Brandes *et. al.* Cys172), and Ssz1 (“yeast OxiAge” Cys sites 86 and Brandes *et. al.* Cys 81). Commonly identified cysteine residues that grouped in clusters A-D and exhibit an increase in oxidation during ageing in both datasets is shown in Figure 6I.

Next, we compared proteins with reactive thiols between different species. We mapped redox-sensitive proteins oxidized during different phases of ageing in wild-type nematode *C. elegans* (Knoefler et al., 2012), wild-type fruit flies (Menger et al., 2015), and young and old mice (Xiao et al., 2020) to the “yeast OxiAge” dataset. We used around 130 proteins from the worm dataset with identified oxidation states. We found 21 proteostasis-regulating proteins common with the yeast orthologs found oxidized in our proteostasis subset (Figure 6D and Supplementary Figure 6C). The small list of common proteins included four found in clusters A-D, namely Eft-3 (yeast Tef1), Dars-1 (yeast Dps1), Cct-8, and Cct-2 (two peptides found in different clusters, B and D) (Figure 6D). Oxidation of 9 evolutionary conserved cysteine residues between yeast and *C. elegans* was identified but none of them belonged to the early oxidation targets grouped in clusters A-D (Figure 6E and Supplementary Figure 6D, Supplementary Table 8). Similarly, the rather small fruit fly dataset (121 proteins and 252 cysteine residues after filtering; see the Methods section) contained only two proteins that were found early oxidized in the “OxiAge” dataset: CNBP (Gis2 in yeast) and Prx1/2/4 (Tsa1 in yeast) (Figure 6F). In total, six common proteins were identified across different clusters with no commonly oxidized cysteine residue (Supplementary Table 8).

A visibly larger number of proteostasis-regulating proteins was commonly found between mice and yeast. On average, ∼50% of yeast proteins had homologs quantified in the mice study (Figure 6G, Supplementary Figure 6E, Supplementary Table 8). Among proteins oxidized during yeast early and mid-point ageing (clusters A-D), the highest proportion of common proteins (∼65%) was seen in cluster B (Figure 6G). The mice study contained data obtained from only two time points, 16-week-old young and 80-week-old mice, but it allowed for unique tissue-specific identification of redox-sensitive proteins. We observed that the percentage of quantified proteins common with yeast proteostasis sub-dataset is comparable between tissues (Supplementary Figure 6G). The highest number of common early oxidized proteins (clusters A-C) is found in brain tissue (19 proteins, >75% of cluster B), followed by brown adipose tissue (BAT), epididymal fat (epi), kidney, liver, subcutaneous fat (subQ), and spleen (17 proteins each, >65% of cluster B). We identified 185 evolutionary conserved redox-sensitive cysteine sites between mice and yeast belonging to proteins within the proteostasis sub-dataset (Figure 6H and Supplementary Figure 6F). Among them, 66 were within the early oxidation targets in cluster A-D. The total number of common cysteine residues was spread unevenly between tissues (Supplementary Figure 6H, Supplementary Table 9). Clusters A and D exhibited the smallest number of common cysteine residues quantified in subQ and liver, respectively, while the highest number of 12 was identified in BAT for early oxidized cysteine residues (cluster A). For 16 proteins with Cys-peptides from clusters A-D in different tissues, we observed an increase in oxidation of more than 5% between young and old mice (Figure 6J), while for an additional 15 an increase of less than 5% (Supplementary Table 8). Among the proteins with evolutionarily conserved cysteine residues showing an age-dependent increase in oxidation in mice were proteins involved in translation. This includes ribosomal proteins [e. g. Rpl34 (Sc_Rpl34A/B), Rps26 (Sc_Rps26B)], ribosome-associated factors [e.g. Eif2S3 (Sc_Gdc1), Eif2S2 (Sc_Sui3), Eif5 (Sc_Tif5), Cnbp (Sc_Gis2)] and tRNA synthetases [e.g. Tars1 (Sc_Ths1)]. Additionally, common proteins with increased % oxidation during ageing included the ubiquitin ligase Znf598 (Sc_Hel2), ubiquitin protease Mocs3 (Sc_Uba4) and components of the TRiC/CCT chaperone complex [e.g. Cct55 (Sc_Sct5), Cct77 (Sc_Cct7)] and cytosolic co-chaperone Dnaja2 (Sc_Ydj1) (Figure 6J).

The integration of the newly described “yeast OxiAge” dataset with existing datasets in yeast, worms, fruit flies, and mice provides the opportunity to identify evolutionarily conserved cysteine residues that are redox-sensitive and to study the dynamics of oxidative modification during ageing. Furthermore, it allows the targeted analysis of proteins and cysteine residues that were not experimentally identified in each organism, and in this respect, the existing datasets complement the “yeast OxiAge” dataset.

### An online tool that allows easy search and visualization of cross-species comparison of proteome oxidation during ageing

To provide an easily accessible interphase for the research community we developed the first cross-species database, called OxiAge Database, that can be accessed through a web application (http://oxiage.ibb.waw.pl; Figure 7). The database contains information on the oxidation levels of ageing yeast (this work and (Brandes et al., 2013)), worms (Knoefler et al., 2012), fruit flies (Menger et al., 2015), and mice (Xiao et al., 2020). Our compendium contains information on the oxidation of 13,051 proteins in canonical or isoform forms (3,153 in yeast, 138 in worms, 420 in flies, 9,340 in mice) and a total of 43,440 cysteine-containing peptides. Each peptide of currently available UniProt ID and detected in any of the datasets has a record in the database. The results are unfiltered, allowing the users to filter according to their interests. Additionally, the user can download the filtered “yeast OxiAge” dataset with the basic analysis presented in this work. The number of records containing information about oxidation levels at different time points/ samples is above 711,000 (Supplementary Figure 7). The web application allows for comparison between proteins with detected cysteine residues and proteins that were not found in any of the datasets but their genes are known orthologs of the detected ones. Thus, the total number of all proteins included in the database reaches almost 26,000 for four different species. The number of ortholog pairs between them is above 76,100.

**Figure 7.**
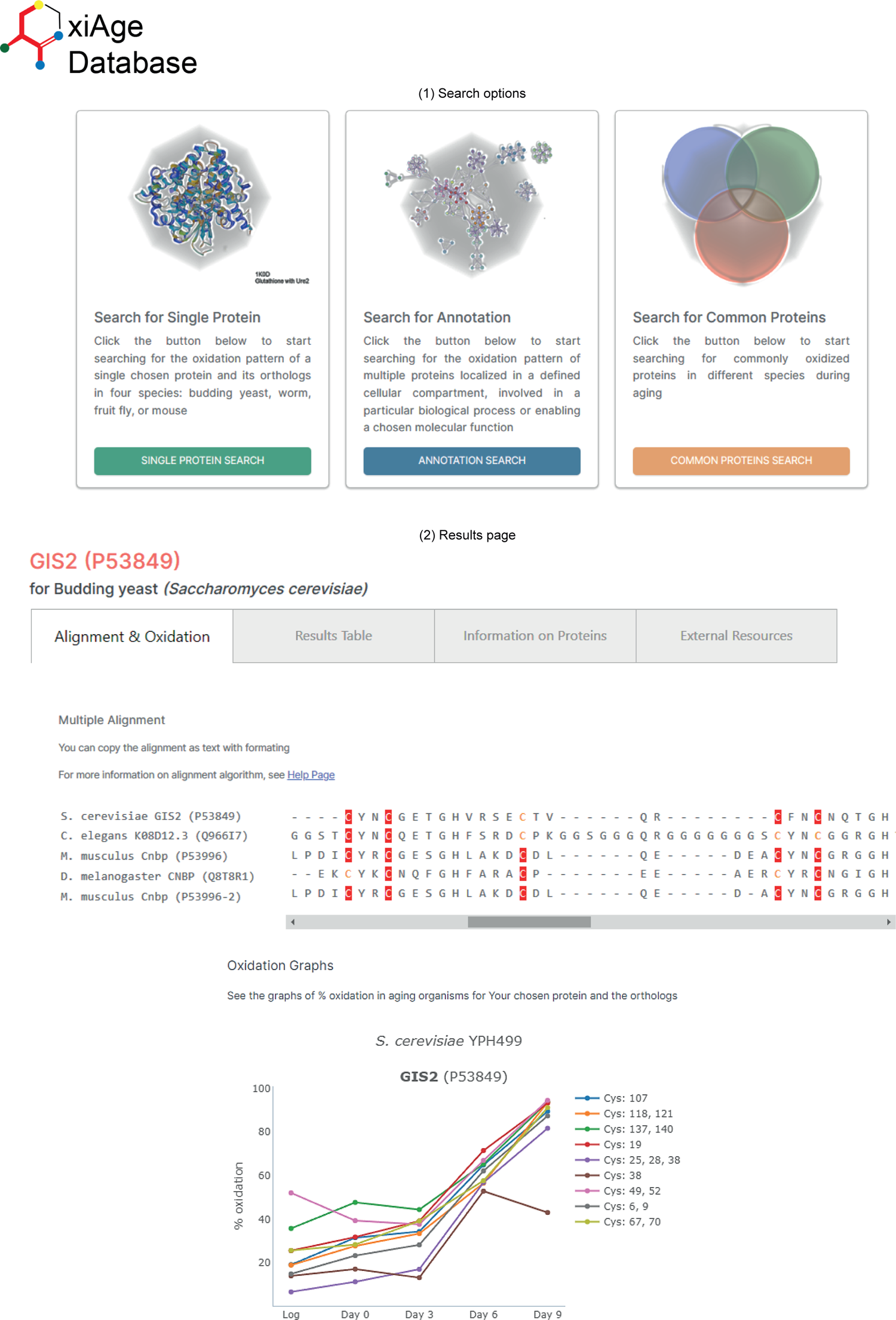
Design of the OxiAge web application. Screenshots of OxiAge Database web application. (1) Search options: The user can choose three options for searching the database: for individual protein/gene name, multiple redox-sensitive proteins common between different species, or for proteins annotated in GO term of chosen biological process, molecular function or cellular component. (2) Results page: The final output of the search presents detailed information about cysteine residues of multiple species found oxidized during ageing. The final results are presented in the form of three tabs. Within the first tab, amino acid alignment is shown with detected cysteines marked in red (information on cysteine position is visible in a tooltip). Graphs present the average % oxidation of cysteines within each peptide. Each found protein and species have its separate graph. Graphs for multiple mouse tissues are presented at the bottom of the tab. Details about the particular experiment and tables containing detailed information about oxidation are shown next to each graph. The second tab contains a table with collected oxidation data for all species and experiments from tab one. Within the third tab, detailed information on chosen proteins and orthologs is found along with the table containing motifs surrounding all cysteine residues. Within the fourth tab, links to external resources for chosen proteins and orthologs are found.

The database allows three distinct types of searches (Figure 7), including searching for a single specified protein within a chosen organism or searching for multiple proteins within a specific biological process, molecular function, or cellular location (according to Gene Ontology annotations). Furthermore, the user can search for all proteins commonly oxidized between two, three or four chosen species. The database is designed to allow easy adjustments for new search procedures and the insertion of additional datasets or information from the existing records (Supplementary Figure 7).

During the search, the user receives information about the searched protein and the orthologs found in other species, along with the sequence alignment with marked cysteine residues detected in a particular organism (Figure 7). Additionally, the user receives results of multiple alignments between proteins with detected cysteine oxidation and orthologs that were not detected within the analysed datasets. Such comparison may serve as a prediction tool for thiol oxidation of proteins that were not found in any of the ageing datasets but may be potential targets of oxidation. The graphs of oxidation levels over time allow for quick determination of the oxidation change for a particular organism and mice tissue (Figure 7). Users can download information about orthologs and protein oxidation. Additionally, a table with all positions of cysteine residues found within a set of proteins, as well as surrounding motifs, is supplied. The results tabs present additional information about searched proteins and orthologs, among them direct links to external resources, such as organism-specific databases (*Saccharomyces* Genome Database, SGD; WormBase: Nematode Information Resource, WB; FlyBase, FB; Mouse Genome-Informatics, MGI), STRING interaction networks, as well as predicted protein structure by AlphaFold (Jumper et al., 2021) or solved crystal structures from Protein Data Bank (PDB).

## Discussion

This study presents a comprehensive dataset on age-dependent protein thiol modifications, providing a rich resource for the research community. The “yeast OxiAge” is accompanied by the OxiAge Database, which is available as a free online tool for the scientific community. The OxiAge Database enables visualization of the results and cross-species comparison of the conservation of oxidation-sensitive cysteine residues. Different search modes enable a targeted approach or allow exploring open-ended scientific questions. The database will be regularly maintained and upgraded. The design of the database and the web application allows for the integration of new datasets that might be published in future, as well as the addition of new search and display functions.

The advantage of the “yeast OxiAge” dataset presented is the temporal resolution of oxidation of 3,178 cysteine residue-containing peptides. Many quantified Cys-peptides enable the unbiased search for early oxidation targets during ageing. Interestingly, our analysis revealed a hierarchy of cellular responses affected by a reversible redox change. At the earliest stage of chronological ageing proteins involved in cellular oxidant detoxification and response to ROS are affected, which is consistent with the observation of a global increase in ROS upon entry into the stationary phase through a TOR-dependent pathway regulating mitochondrial respiration (Pan et al., 2011). This first response of the cell is accompanied by a switch of activity of transcription factors between the logarithmic and stationary phases. These processes are all represented by proteins oxidized during the entry into the stationary phase on day 0 and are mostly associated with oxidation cluster C. The following processes are involved in global proteome homeostasis through changes in protein production (cluster A) for which we observed the regulation of ribosome assembly and cytosolic translation. Moreover, there is a visible change in the oxidation of subunits of polymerase I required for the transcription of rRNA. This indicates that cells prioritize the regulation of global protein synthesis above other processes during the very early stages of ageing possibly adapting quickly to the increasing protein burden in the cell. The regulation of translation is a common stress response and a decrease in global protein production was shown to be beneficial for lifespan in model organisms when targeting components of the translation machinery early in their lives (Anisimova et al., 2018; Chiocchetti et al., 2007; Curran & Ruvkun, 2007; Hansen et al., 2007; Rogers et al., 2011; Steffen et al., 2008; Tavernarakis, 2008; Topf et al., 2019; von der Haar et al., 2017). Prominent in our analysis was the effect on ribosomal proteins, which also was identified during stress conditions in previous studies (Kumsta et al., 2011; Marino et al., 2010; Menger et al., 2015; Topf et al., 2018). Still, a mechanistic consequence of the reversible oxidation of ribosomal proteins remains to be elucidated. Moreover, we identified Gis2, a translational activator for mRNAs with internal ribosome entry sites, to have several conserved cysteine residues sensitive to oxidation. The role of Gis2 during ageing is unknown but it was shown previously that Gis2 and its human homolog CNBP are associated with stress-induced RNP granules suggesting a role in translational repression (Rojas et al., 2012). Moreover, several tRNA synthetases were among the targets of early oxidation. Our observation confirms previous findings in mice that identified the tRNA synthetase proteins as the most ubiquitously oxidized between different tissues in comparison to other proteins (Xiao et al., 2020).

Increasing protein burden through increased levels of ROS in the cell during ageing triggers further adaptation in cellular processes including changes in the global transcription of mRNA through regulation of polymerase II and protein degradation (cluster B). We found that regulation of protein degradation rather follows the adaptation of translation and transcription. Here, enzymes of the ubiquitination cycle were affected. The sensitivity of the active site cysteine residues of ubiquitinating and deubiquitinating enzymes was shown previously. For example, oxidized glutathione was shown to react with the active site cysteine in E1 and E2 forming mixed-disulfide bonds and blocking their binding to ubiquitin (Jahngen-Hodge et al., 1997). However, the activity of deubiquitinating enzymes (DUBs) assures a balance in the ubiquitinated proteome. DUBs are especially sensitive to oxidative stress. A single molecule, such as hydrogen peroxide, can affect a vast number of DUBs without the need for an enzymatic intermediate (Snyder & Silva, 2021). Further, functional links between ubiquitination factors and translation were established (Kapadia & Gartenhaus, 2019). Protein synthesis can be controlled by the ubiquitination of ribosomes. Oxidative stress induces high levels of K63-linked polyubiquitin chains on ribosomes (Dougherty et al., 2020). Polyubiquitination pauses translation elongation (Zhou et al., 2020). Interestingly, we found that the ubiquitin-specific protease Ubp1 was early oxidized during ageing. Ubp1 was recently found to form a disulfide bond with the ubiquitin ligase Rad6, which is a determinant of ubiquitinated ribosomes. This reversible thiol modification between Ubp1 and Rad6 inhibited K63-linked ubiquitin modification (Simoes et al., 2022). A possible implication of this mechanism during ageing remains to be discovered.

As cells get older and enter the middle stage of ageing, a group of folding chaperones (cluster D) is increasingly oxidized. Here, protein chaperones downstream of the protein synthesis machinery seem affected. Chaperones of the Hsp70 family in general, including Ssz1 and Ssb1 identified in this study, were shown previously to act as redox sensors (Zhang et al., 2022). While often the specific physiological consequences of cysteine residue modification of particular Hsp70s are unknown, their oxidation can change the fate of their targets thereby amplifying redox signalling to a broader range of proteins achieving a greater antioxidative potential (Zhang et al., 2022). This includes the increased expression of chaperones. Newly, synthesized Hsp70 chaperones are not oxidized and can prevent protein aggregation and/or facilitate the degradation of oxidized proteins (Kalmar & Greensmith, 2009). We also found that subunits of the TRiC/CCT complex showed increased oxidation on evolutionarily conserved cysteine residues. The TRiC/CCT complex is an ATP-dependent chaperonin responsible for folding of a variety of cellular proteins (Gestaut et al., 2019). Its potential regulation by oxidation is unknown but due to its high abundance in the cell might also act as a cellular redox sensor. Thus, our analysis provides a rich resource of redox-sensitive key proteins during the early phase of yeast ageing governing processes of the protein homeostasis network.

While our study is limited to the mere identification of oxidative thiol modifications and their dynamics during ageing, for many examples we can only speculate about the role of the shift in the redox state. Most importantly, following studies will need to address the question of whether the increase in the oxidized pool of a particular protein acts as a stress response aiming to rescue detrimental effects by restoring protein homeostasis during ageing or is a driving force of the ageing process.

## Methods

### Yeast growth conditions

For OxICAT analysis, *Saccharomyces cerevisiae* strain YPH499 (*MATa, ade2-101, his3-Δ200, leu2-Δ1, ura3-52, trp1-Δ63, lys2-801*) was grown in three biological replicates at 28°C on minimal synthetic medium (0.67% [w/v] yeast nitrogen base, 0.079% [w/v] CSM amino acid mix, 20 mg/l [w/v] Adenine) containing 2% [v/v] glucose. Samples were harvested by centrifugation (3500x g, 5 min) at the log phase (OD_600_ ∼0.45), when they had reached stationary phase (=Ageing Day 0, OD_600_ ∼7) and at days three, six and nine during stationary phase. Pellets from 50 ml culture (Log phase) and 10 ml culture (day 0, 3, 6, 9) were resuspended in 10% TCA, 150 mM NaCl, frozen in liquid nitrogen and stored at −80°C

### Yeast survival assay

To determine yeast viability, one OD600 unit from each culture of chronologically aged yeast was collected by centrifugation at 3000 × g for 5 min at room temperature. Pellets were resuspended in 1 ml of 1x PBS (137 mM NaCl, 12 mM KPO_4_, 2.7 mM KCl, pH 7.4). Propidium iodide (Sigma-Aldrich, catalogue no. P4170, 3 µg/ml final concentration) was added to the cell suspension and incubated for 15 min at room temperature while protected from light. Propidium iodide stains only dead cells. A negative control (no staining with propidium iodide) and a positive control were prepared for each sample. For the positive control yeast cells were heat-killed by incubation at 80°C for 2 min. Samples were diluted 1:10 in 1x PBS and kept on ice and analyzed within one hour after staining. All samples were analysed by flow cytometry (BD FACS CALIBUR). 30000 cells per sample were analysed. Number of dead cells was recorded using BD CellQuest Pro software.

### OxICAT labelling

Protein extraction from whole cells, differential thiol trapping, tryptic digest, peptide enrichment and cleavage of the biotin tag were performed exactly as described before (Topf et al., 2018). Briefly, samples were homogenised in 10% TCA using glass beads. Protein concentration was determined by Bradford assay. 200 µg aliquots were centrifuged (21,000 x *g*, 15 min, 4°C), washed with 5% ice-cold TCA and resuspended in a mixture of 160 µl denaturing alkylation buffer (DAB, 6 M urea, 200 mM tris(hydroxymethyl)aminomethane-HCl [Tris-HCl], pH 8.5, 10 mM ethylenediaminetetraacetic acid [EDTA], 0.5% SDS) with 2 units of heavy ICAT reagent (ABSciex) dissolved in 40 µl acetonitrile. Samples were incubated for 2 h at 37°C under vigorous shaking. Proteins were precipitated by acetone and resuspended in a mixture of 160 µl DAB with 2 units of light ICAT reagent dissolved in 40 µl acetonitrile. TCEP was added to a final concentration of 1 mM and samples were incubated for 2 h at 37°C under vigorous shaking followed by acetone precipitation. Protein pellets were digested in 200 µl of 20 mM ammonium bicarbonate (NH_4_HCO_3_). with 12 µg Trypsin (sequencing-grade modified, Promega). After 16h incubation at 37°C insoluble proteins were pelleted by centrifugation (21,000 x *g*, 1 min, room temperature) resuspended in 50 µl of 60% (v/v) methanol, 20 mM NH_4_HCO_3_ and incubated with 1 µg Trypsin at 42°C for 3 h. Supernatants of digests in 20 mM NH_4_HCO_3_ and in 60% (v/v) methanol, 20 mM NH_4_HCO_3_ were combined. Clean-up of the digests by strong cation exchange (SCX) chromatography, enrichment by avidin affinity chromatography and cleavage of the biotin tag were performed using the ICAT kit (AbSciex) following the manufacturers’ instructions.

### LC-MS analysis

Nano-HPLC-ESI-MS/MS analyses were performed at an Orbitrap Elite mass spectrometer connected to an UltiMate 3000 RSLCnano HPLC system (Thermo Scientific) exactly as described before for whole cell oxICAT samples (Topf et al., 2018). Peptide samples were resuspended in 30 µl 0.1% TFA, and each biological replicate was analysed in two technical replicates.

### Mass spectrometry data analysis

Raw files of the LC-MS/MS runs were jointly processed using MaxQuant v 2.0.2.0 (Cox & Mann, 2008). Peptides corresponding to the *Saccharomyces cerevisiae* protein database (UniProtKB canonical set for strain ATCC 204508 / S288c, Proteome ID UP000002311, were retrieved on May 2^nd^, 2022, 6091 entries) using Andromeda (Cox et al., 2011). ICAT light tag (C_10_H_17_N_3_O_3_, 227.13 Da) and ICAT heavy tag (^13^C_9_CH_17_N_3_O_3_, 236.16 Da) were set as light and heavy labels, respectively, with cysteine residue specificity. Up to five labelled amino acids per peptide were allowed. Trypsin was set for the generation of theoretical peptides, allowing a maximum of two missed cleavages. Oxidation of methionine was included as a variable modification. Labelling ratio estimation was performed using the options “Re-quantify” and “Match between runs” with a retention time window of 0.7 min. The precursor mass tolerance was set to 20 ppm for the “first search” option of Andromeda and to 4.5 ppm for the main search. The minimum peptide length was set to 6 amino acids. A peptide spectrum match (PSM) false discovery rate (FDR) of 1% was applied using the decoy mode “Revert”.

The MaxQuant file “evidence.txt” was used for subsequent analysis. Entries from potential contaminants, decoy hits and evidences with no quantified intensity were removed. Intensities for the heavy or light labelled peptides of 0, resulting from MaxQuant identifying but not quantifying a peptide, were set to NA. Peptide starting positions (i.e., the sequence index of the first amino acid of the peptide) were extracted from the MaxQuant file “peptides.txt” and used for calculating absolute positions. Cysteine identifications (CysIDs) for all peptides containing a cysteine residue were generated from the protein identifier of the leading razor protein and the absolute position of the cysteine included in the peptide. Peptides with multiple cysteine residues were assigned a CysID including all identified residues. Next, % light intensity was calculated for each evidence by dividing the intensity of light-labelled peptides by the sum of intensities of light and heavy-labelled peptides. ISO-MSMS evidences with labelling state 0 (i.e. only the light labelling partner was identified) which exhibited % light intensities below 50% after re-quantification were removed. Likewise, ISO-MSMS evidences with labelling state 1 (i.e. only the heavy labelling partner was identified) which exhibited % light intensities above 50% after re-quantification were removed. MULTI-MATCH and MULTI-SECPEP evidences were filtered out if each ISO-MSMS evidence identified for the cysteine residue (i.e. miscleaved, methionine oxidation, different *z* values) was observed with labelling state 0 and % light intensity < 50% or respectively labelling state 1 and % light intensity > 50%. The resulting table contained 224469 entries (corresponding to 97.2 % of all identified cysteine-containing evidences with non-zero intensity) and was used to calculate the proportion of reversibly oxidized cysteine residues, which we refer to as % oxidation, for each unique cysteine residue. For each timepoint and biological replicate total intensities (i.e. light + heavy) and light intensities respectively of both technical replicates were summed up over all evidences (i.e. modified, miscleaved, different charge states) of a given CysID. The % oxidation was calculated as the sum of light intensity divided by the sum of total intensity (light + heavy) multiplied by 100 (for a detailed description, see (Topf et al., 2018)).

All raw values corresponding to day 3, biological replicate 3, were removed as this replicate showed disproportionately higher missing values (27% for light-labelled and 33 % for heavy-labelled CysID evidences) compared to the other replicates (< 5% for light- and heavy-labelled CysID evidences, respectively). Furthermore, CysIDs with missing heavy or light intensities in more than one biological replicate were removed. In total, 65.5% of all peptides were removed after filtering. 3178 peptides of a total of 9213 remained for further downstream analysis. We refer to this dataset as “yeast OxiAge” dataset.

### Missing values imputation

Missing intensity values for light or heavy ICAT-labelled peptides resulting from unidentified or non-quantified peptides (see data processing) were imputed using a data-driven imputation strategy (Egert et al., 2021). The filtered data frame was used as input for the DIMA protocol implemented in R (version 4.1.0). Briefly, DIMA uses a subset of the data frame with no missing values as a template to generate a benchmarking dataset with the same distribution of missing values as the original data set. Various imputation algorithms are tested on the benchmark dataset and evaluated based on the known true values. The best-scoring algorithm is used for the imputation of the initial dataset. For the OxICAT dataset, the impSeqRob algorithm from the R package rrcovNA (Branden & Verboven, 2009; Verboven et al., 2007) was consistently best-performing. Using this imputation method, a total of 2345 intensity values for light and heavy ICAT-labelled peptides were generated (2.64% of all intensity values), corresponding to 1229 values of % oxidation (2.76% of all oxidation values). Dataset with imputed values was used to generate principal component analysis (PCA) of components 1 and 2 for each replicate and each time point and volcano plots of statistically significant changes between each pair of time points.

### Peptides clustering

All downstream analyses of the filtered mass spectrometry dataset were performed in Python version 3.9 unless stated otherwise. Filtered “yeast OxiAge” dataset with not imputed values was used for downstream analysis. Peptides were clustered into ten groups based on the mean change in % of oxidation over time in a two-step process. Firstly, outliers that could impact the clustering procedure were determined using the DBSCAN algorithm implemented in the Python package sklearn (https://github.com/scikit-learn/scikit-learn, (Pedregosa et al., 2011)) with a maximum distance between neighbouring samples set to 22 and the total weight set to 13. Six outliers were identified and removed from the clustering dataset. Secondly, peptides after the removal of the outliers were grouped using Standard Scaler for segmentation followed by Gaussian Mixture (GM) for clustering, both algorithms implemented in the sklearn package. Peptides were grouped into four initial clusters, as determined by the Elbow method, with random seed set to 8000, convergence threshold set to 1 and covariance type set to diag (diagonal covariance matrix per component). Clusters were reviewed manually and standard deviations between clusters were compared. Clusters exhibiting nonhomogeneous trends were re-clustered using the same parameters and the random seed set to 10000, as follows: two similarly behaved clusters 1 and 3 were re-clustered together into 5 clusters; cluster 4 exhibiting the highest standard deviation was re-clustered into 4 separate clusters; cluster 2 was not re-clustered due to small standard deviation within the cluster. The resulting 10 clusters were manually labelled A-J based on the trend of an early increase in % of oxidation. The six outliers removed from the dataset before clustering were manually added to the cluster containing the highest variation between peptides with no obvious trends (cluster I). The method of clustering, the number of clusters used and the exact parameters’ values were decided by the try-and-check method.

### Annotation analysis

Gene Ontology (GO) annotations for cellular components (CC) of yeast proteome were retrieved on 14.03.2022 from UniProtKB for strain ATCC 204508 / S288c, Proteome ID UP000002311. For cellular localization, GO annotations were consolidated into nine main annotations according to parent-child terms, using as the highest degree parent terms corresponding to Cytoplasm (GO:0005737), Nucleus (GO:0005634), Endoplasmic reticulum (ER, GO:0005783), Golgi apparatus (GO:0005794), Mitochondrion (GO:0005739), Vacuole (GO:0005773), Peroxisome (GO:0005777; GO:0019818), Lysosome (GO:0005764), and Ribosome (GO:0005840; GO:0033279). An additional group named “Other” contained all proteins with other localizations specified that were not considered in this study, such as extracellular region, etc. Proteins that belong to both terms Cytoplasm and Nucleus were grouped into a separate term “Nucleus / Cytoplasm”. Proteins associated with cytosolic and mitochondrial ribosomes were grouped into the term “Ribosome”. Proteins associated with Cytoplasm and a second additional organelle except for Nucleus, were considered annotated to the latter group, i.e. a protein assigned to Cytoplasm and ER was grouped as ER-localised. Proteins assigned to multiple organelles were grouped under the term “Multiple localization”. Proteins without a GO annotation specified were grouped under the term “Unknown”.

To identify proteins involved in proteostasis, we used GO Slim Yeast annotations retrieved from QuickGO (Binns et al., 2009) on 27.10.2022. We grouped proteins according to one of three groups of biological processes (BP) regulating proteostasis: cytoplasmic translation, protein degradation, or protein folding. For global translation, we searched for the words “translation”, “translational” etc. (exception: mitochondrial-specific translation). For protein degradation, we searched for the words “ubiquitin”, “ubiquitination”, “polyubiquitination”, “proteasome”, “proteasomal”, “protein conjugation”, and “proteolysis”.

For protein folding, we looked for the words “folding” and “refolding”. Results were manually reviewed for uncertain proteins. Proteins within each cluster (A-J) were grouped based on one of the three proteostasis groups.

Information about disulphide bonds, protein domains, length, sequence, isomers, structures, interactions, etc. was retrieved from UniProtKB from 14.11.2022 – 03.02.2023.

### GO enrichment analysis

Analysis of the overrepresentation of GO terms was performed using the TopGo package in R (version 4.1.0). For each cluster, a custom analysis was performed using as an input species yeast *S. cerevisiae* and as a background cysteine-containing proteins according to amino acid sequences provided from the yeast proteome (yeast proteome retrieved on 14.03.2022 from UniProtKB for strain ATCC 204508 / S288c, Proteome ID UP000002311). We identified 5478 canonical proteins as containing at least one cysteine residue within the provided amino acid sequence. GO Slim annotations for yeast were retrieved from QuickGO (Binns et al., 2009) on 27.10.2022. GO Biological Process (BP) and GO Molecular Function (MF) were searched for enrichment. GO terms with at least 20 genes per term were used and the algorithm weight01 with Fisher statistics was used to calculate the p-values for each term. The top 200 terms were selected and *p*-value correction was performed on GO terms with fold enrichment > 1.5 and *p*-value < 0.05. Benjamini-Hochberg’s (BH) procedure was used as a correction for multiple testing for the identification of significant enrichment. Additionally, Bonferroni and Hochberg corrections were performed. The fold enrichment was calculated as the number of significant genes found in a GO term divided by the number of expected genes taken from the submitted background. Fold enrichment is a measure that indicates how drastically proteins belonging to a certain pathway are overrepresented independently of the pathway size. To simplify and summarize the high number of enriched processes, the BP terms were grouped using the online tool Revigo (Supek et al., 2011), with the GO database updated on 03.2022 and UniProt-to-GO mapping from EBI GOA updated on 04.2022. *S. cerevisiae* was used as a species, the Lin method was used as a semantic similarity measure and C-value was set to 0.5. For visualization purposes, a representative GO term for each Revigo group was chosen based on the lowest *p*-value.

### Identification of zinc-binding cysteine residues

To determine zinc-binding proteins, the dataset established by Wang et al. (Wang et al., 2018) was used. All oxidized cysteine residues per each quantified protein from the “yeast OxiAge” dataset were mapped into known or potential zinc-binding proteins (582 proteins) and zinc-dependent proteins identified by mass spectrometry analysis of protein abundance in zinc-depleted cells (over 2500 proteins identified).

### Protein association network

Cytoscape 3.9.1.(Shannon et al., 2003) with a plug-in StringApp 1.7.1. (Doncheva et al., 2019) was used to generate a functional protein association network. The minimum required interaction score was set to confidence 0.6 unless specified differently. A full STRING network was applied with the meaning of network edges set to “confidence”, meaning that the thickness of lines indicates the strength of evidence in the literature. For the display, the structure previews were disabled. A compound spring embedder layout was used for visualization. Proteins were coloured based on the biological process of proteostasis: translation in blue, folding in green, and/or degradation in red. Proteins that belong to more than one group had nodes coloured with more than one colour.

### Motif analysis

Analysis of motifs was performed by extracting motifs centred ±6 amino acids around each quantified cysteine residue in the “yeast OxiAge” dataset. The analysis and visualization of motifs with a total length of 13 amino acids were performed using the freely available online tool pLogo (O’Shea et al., 2013). Overrepresented and underrepresented motifs were analysed for each A-J cluster separately. Cysteine residues detected in more than one peptide were used only once. Sequences with cysteine residues closer to the START or STOP codons than 6 amino acids were removed from the analysis. As background, sequences in the FASTA format of yeast proteins containing cysteine residues (obtained from yeast proteome in UniProtKB as described before) were used. Redundancy was removed from the background and the foreground. Forward sequences were not removed from the background. Amino acids with *p*-values < 0.05 were considered significant. Colours were assigned to each amino acid: nonpolar amino acids are in black except G and A coloured in green; sulfur-containing methionine (M) is in black; polar uncharged are in orange; aromatic are in grey; positively charged are in red; negatively charged are in blue; sulfur-containing cysteine (C) is in yellow. Log-odds of binominal probability are shown and frequencies of each overrepresented amino acid were extracted.

### Ortholog analysis

To compare data between different species, the orthologs file was downloaded from the Alliance of Genome Resources on 16.05.2022 (Alliance Database Version 5.2.0; species: *Homo sapiens, Rattus norvegicus, Mus musculus, Danio rerio, Drosophila melanogaster, Caenorhabditis elegans, Saccharomyces cerevisiae*). Canonical forms of proteins quantified in the yeast dataset were searched for orthologs in other species based on identification numbers from a specific organismal database: Saccharomyces Genome Database (SGD), Mouse Genome Informatics (MGI), WormBase: Nematode Information Resource (WB), and FlyBase (FB). The data were compared between different species of datasets: “yeast OxiAge” (this study), yeast strain DBY746 (Brandes et al., 2013), nematode (Knoefler et al., 2012), fruit fly (Menger et al., 2015), and mice (Xiao et al., 2020). The four latter datasets were filtered before the orthologs analysis. For yeast strain DBY746, data were retrieved from the experiment performed in standard conditions for chronological ageing (2% glucose). Values for % of oxidation at the logarithmic phase (Log), days 1, 2, 3, and 4 were obtained. Peptides with unidentified or unquantified average % oxidation values in any of the given time points were removed from the analysis and comparison. UniProt IDs were manually checked and updated to the current state of the database on day 24.10.2022. Obsolete terms were removed. In total, 287 peptides were retrieved. For nematode *C. elegans* strain N2 “Bristol”, data were retrieved from the experiment performed in wild-type animals from the larva stage L4 and worms at days 2, 8 and 15 of adulthood. Filtering of the peptides was performed as described before for yeast strain DBY476. UniProt IDs were manually checked and updated, and obsolete terms were removed, as described before. In total, 108 peptides were retrieved. For fruit flies, data from wild-type days 7, 28 and 56 of adulthood were retrieved for control strains, ageing. We considered only peptides with identified values for three time points (at least three repeats had detected and quantified intensity according to initial filtering performed by *Menger et al.*). 252 peptides were retrieved. For male mice *M. musculus*, strain C57BL/6, data on young and old mice were obtained from the analysis performed on ten different tissues in animals aged 16 and 80 weeks. Filtering of the peptides was performed as described before. UniProt IDs were manually checked and updated, and obsolete terms were removed, as described before, for each time point and tissue. In total, 60841 peptides for different tissues were retrieved, which corresponds to 21850 unique peptides in all tissues. Evolutionarily conserved proteins found oxidized in different species were labelled as “common proteins”.

For comparison of evolutionarily conserved cysteine residues, sequence alignment was performed (see below) between yeast proteins involved in proteostasis and corresponding proteins in worms and mice identified by orthologs search. Positions of the aligned cysteine residues were matched to the positions of the residues of oxidized cysteines within different datasets. Conserved cysteine residues found oxidized in different species were labelled as “common cysteines”.

To visualize the overlap between species, all oxidized proteins in the worm, fruit fly and mouse datasets were labelled according to the yeast orthologs UniProt ID, then mouse and worm, unless no orthologs were found.

### Sequence alignment

Amino acid sequence alignment was performed using Biotite framework version 0.35.0 (Kunzmann & Hamacher, 2018) implemented in Python, Biopython version 1.80 (Cock et al., 2009). Clustal Omega (Sievers & Higgins, 2021) was performed for multiple sequence alignment with a Python package clustalo.ClustalOmegaApp utilizing the software ClustalOmega version 1.2.2. For pairwise alignment packages align_optimal and SubstitutionMatrix.std_protein_matrix from Biotite.align were used. A custom-made code in Python was developed to obtain detailed information about the positions of aligned cysteine residues in different species.

### Database and web application

The web application (http://oxiage.ibb.waw.pl) is currently hosted on the Institute of Biochemistry and Biophysics Xen 6.2 virtualization on Ubuntu 22.04 LTS operating system, Apache 2 version 2.4 wsgi modem. It is developed using Python Interpreter version 3.10 in the Dash framework version 2.7.1 (https://github.com/plotly/dash). For display and user interaction Javascript is used. Cascading Style Sheets (CSS) is used for styling website components. Graphs and charts are displayed using plotly.js (https://github.com/plotly/plotly.js/). For responsive layout Dash bootstrap components version 1.3.0 (https://github.com/facultyai/dash-bootstrap-components), Dash mantine components version 0.11.1 (https://github.com/snehilvj/dash-mantine-components), and Dash HTML components version 2.0.0 (https://github.com/plotly/dash-html-components) are used. To visualize data frames and tables pandas package version 1.5.2 (https://github.com/pandas-dev/pandas) and Dash table components version 5.0.0 (https://github.com/plotly/dash-table) are used. Sequence alignment is performed using Biotite.align from Biotite framework version 0.35.0 (Kunzmann & Hamacher, 2018). The database containing static resources (i.e., protein name, protein sequence, detected cysteine sites) is developed using a structural query language (SQL) and a relational database management system MySQL 8.0. Connection to MySQL database is established using MySQL Connector/Python version 8.0.31 (https://github.com/mysql/mysql-connector-python). For a detailed description of the use and features of the database and the web application, please visit the help page of the website.

### Statistical analysis

Number of sample sizes, replicates and statistical tests were chosen according to published data with comparable methodology. No statistical method was used to predetermine the sample size. For statistical analysis of oxidized peptides in filtered “yeast OxiAge” dataset, imputed values were used. Statistical significance between each time point group was calculated using a two-sided Welch’s *t*-test for an unequal variance for imputed data. Volcano plots were used to visualize the data and present -log10 of *p*-value on the y-axis and the difference in mean % of oxidation between each pair. Principal component analysis (PCA) for components 1 and 2 was generated using MaxQuant/Perseus software version 2.0.9.0 (Tyanova et al., 2016). Pearson’s correlation coefficient between all replicates of imputed and nonimputed data was calculated using the Python package sklearn (https://github.com/scikit-learn/scikit-learn, (Pedregosa et al., 2011)). Statistical analysis of differences in % oxidation of peptides between time points per each cluster A-J was performed using Kruskal-Wallis test with Dunn’s post-hoc method. Bonferroni correction was applied for multiple testing and *p*-values were considered significant if < 0.005. For GO enrichment analysis, Fisher’s exact test was used as described before to identify enriched GO terms. Benjamini-Hochberng (BH) procedure was used to correct for multiple testing and identify significantly enriched GO terms. Significant difference in the overrepresentation of motifs using pLogo was calculated as described in (O’Shea et al., 2013). Statistical significant threshold was calculated using Bonferroni correction. The y-axis denotes the log-odds binomial probability. Residues are stacked from most to least significant.

### Data and code availability

The mass spectrometry proteomics data have been deposited to the ProteomeXchange Consortium (Deutsch et al., 2022) via the PRIDE (Perez-Riverol et al., 2022) partner repository with the dataset identifier PXD042047. The web source code is available via GitHub: https://github.com/katarzynajonak/oxiage. The web server is available at: http://oxiage.ibb.waw.pl. The website is free and open to all users. All other data needed to evaluate the conclusions in the paper are present in the paper and/or the Supplementary Materials.

## Author contributions

UT set up culture conditions for yeast chronological ageing assay. UT and IS prepared samples for mass spectrometry analysis. IS prepared proteomics analysis and analysed raw data together with JB. KJ performed downstream bioinformatics analysis. KJ designed and developed the database and web server application. AC and BW conceived the yeast OxICAT study and supervised the experiment and the analysis of proteomics data. UT conceived the bioinformatics study and supervised the analysis. AC, BW and UT provided funding for the research. KJ and UT wrote the initial manuscript and prepared the figures. JB and BW edited the manuscript. All authors agreed on the final version of the manuscript.

## Acknowledgements / Funding

The OxICAT analysis and data acquisition were established and performed in the laboratories of AC and BW. Work in the AC laboratory was supported by National Science Centre Poland (grant no. 2011/02/B/NZ2/01402). Work in the BW laboratory was supported by the Deutsche Forschungsgemeinschaft (DFG, German Research Foundation) Project ID 403222702/SFB 1381, FOR 2743 (WA1598/6), TRR 130 and Germany’s Excellence Strategy (CIBSS – EXC-2189 – project ID 390939984). Work connected to the bioinformatics analysis in the UT laboratory was funded by National Science Centre Poland (grant no. 2019/34/E/NZ1/00367) and additionally supported by the Institute of Biochemistry and Biophysics PAS Internal Grant for KJ (DEC-MG-4/22-03). KJ is supported by EMBO Postdoctoral Fellowship (grant ALTF 82-2022). We thank members of the Laboratory of Mitochondrial Biogenesis for discussions on data interpretation during the process of the OxICAT data generation. We are grateful to Lukasz Knizewski for the IT support and to Karol Jedrasiak for the help with the database and the website design. We would like to thank the members of the Laboratory of Molecular Basis of Aging and Rejuvenation for testing the web application. The funders had no role in the study design, data collection and analysis, or preparation of the manuscript.

---

**Supplementary Figure 1. (Related to Figure 1).**
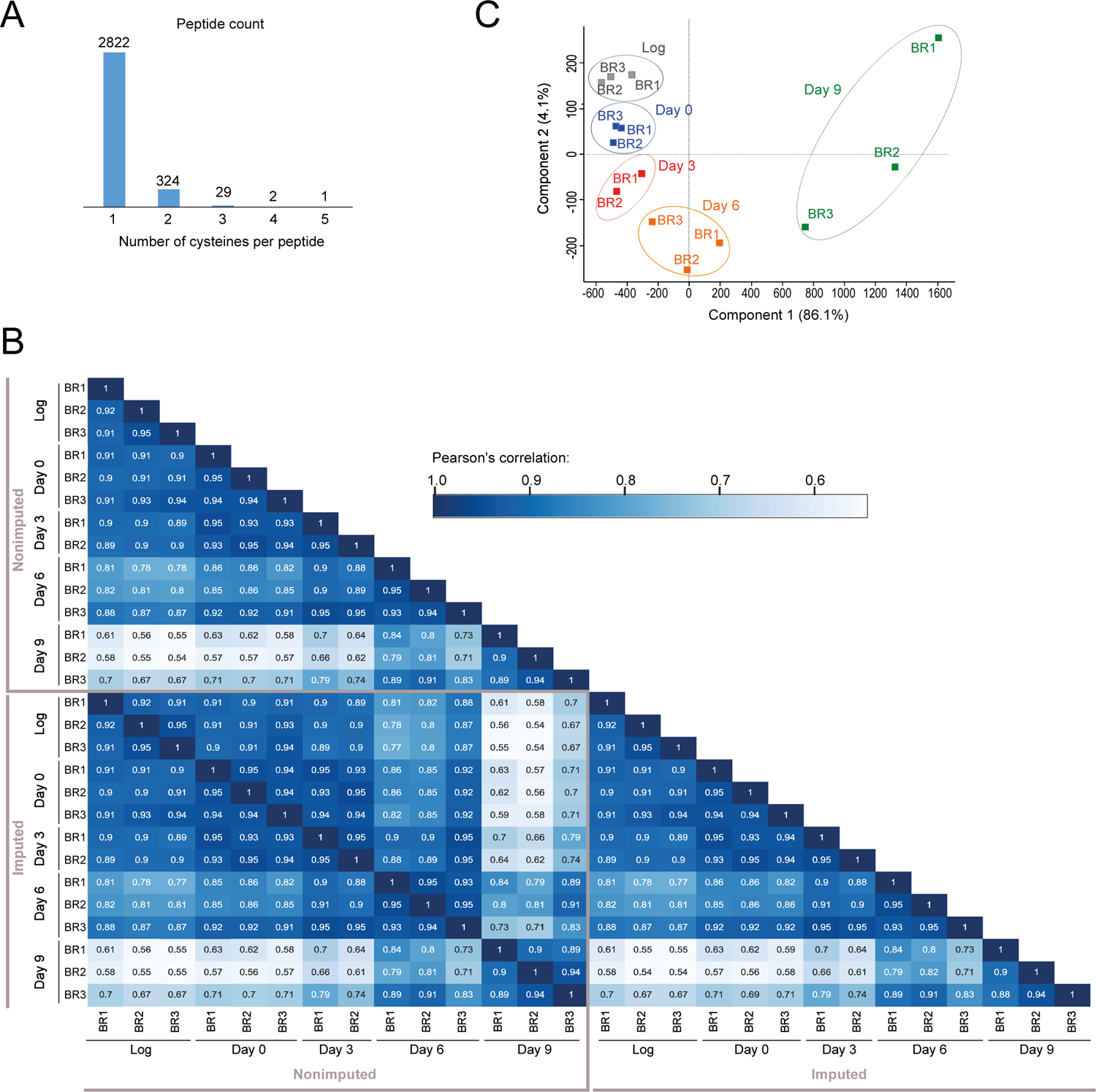
Analysis of the “yeast OxiAge” dataset. **(A)** Number of cysteine residues found within a quantified Cys-peptide. The majority of peptides (>2800) contain only one redox-sensitive cysteine detected and quantified. **(B)** Heatmap of Pearson’s correlation between different biological replicates and time points for not imputed (“Non-imputed”) and imputed (“Imputed”) values. *BR*, biological replicate; *Log*, logarithmic phase. **(C)** Principal component analysis (PCA) plot showing variation between individual biological replicates and time points for the “yeast OxiAge” dataset with imputed values. *BR*, biological replicate; *Log*, logarithmic phase.

**Supplementary Figure 2. (Related to Figure 2).**
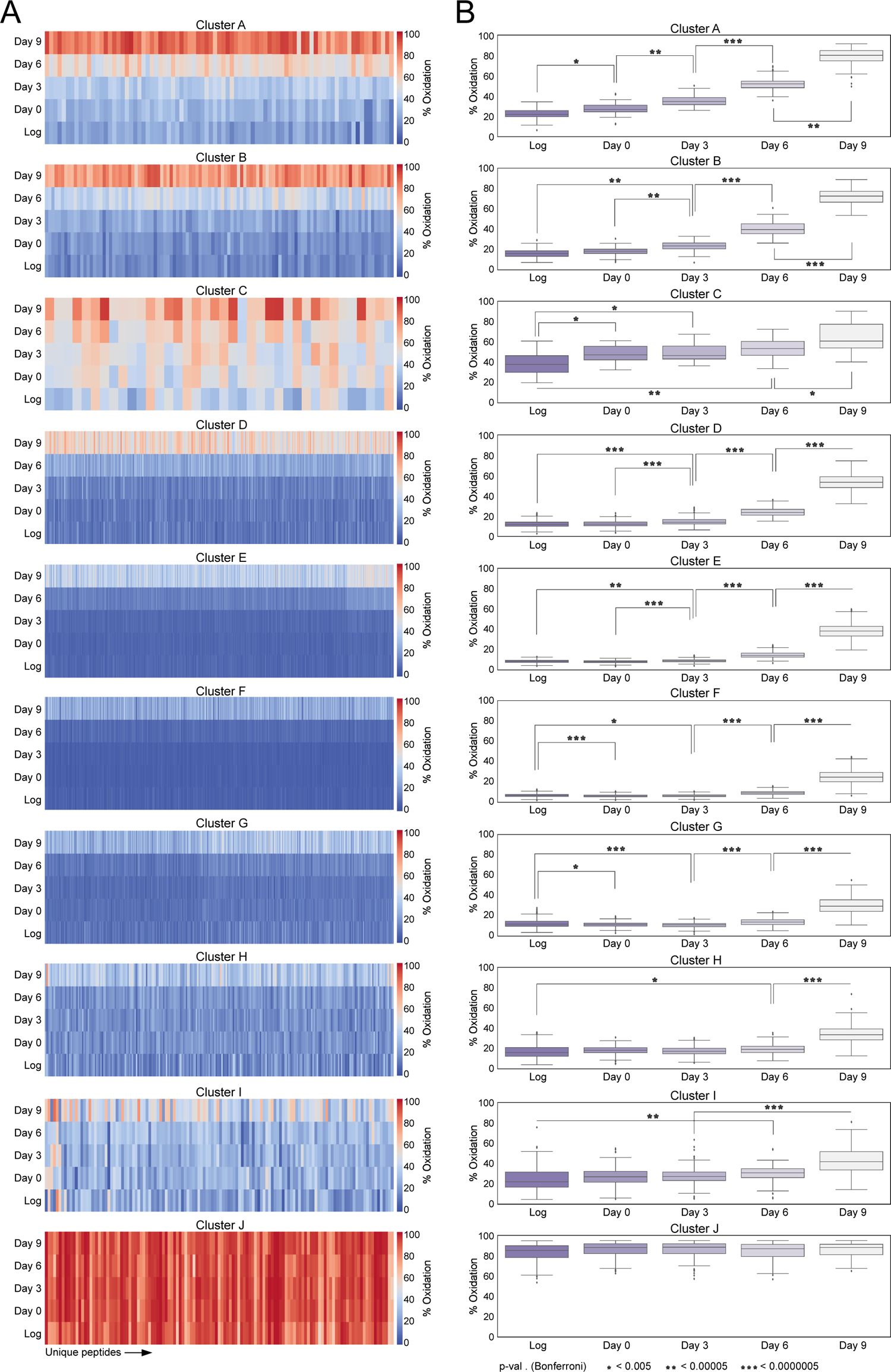
Distinct patterns of sequential peptides oxidation for individual clusters. **(A)** Heatmap of average % oxidation per Cys-peptide on each day for each cluster, A-J. *Log*, logarithmic phase. **(B)** Boxplots of average % oxidation per Cys-peptide on each day for each cluster, A-J. Statistical significance of the difference in % oxidation between clusters was assessed using the Kruskal-Wallis test with Dunn’s posthoc method with Bonferroni correction. Stars indicate the significance level: one, *p*-value < 0.005 (significance threshold for Bonferroni correction); two, *p*-value < 0.00005; three, *p*-value < 0.0000005. *Log*, logarithmic phase.

**Supplementary Figure 3. (Related to Figure 4).**
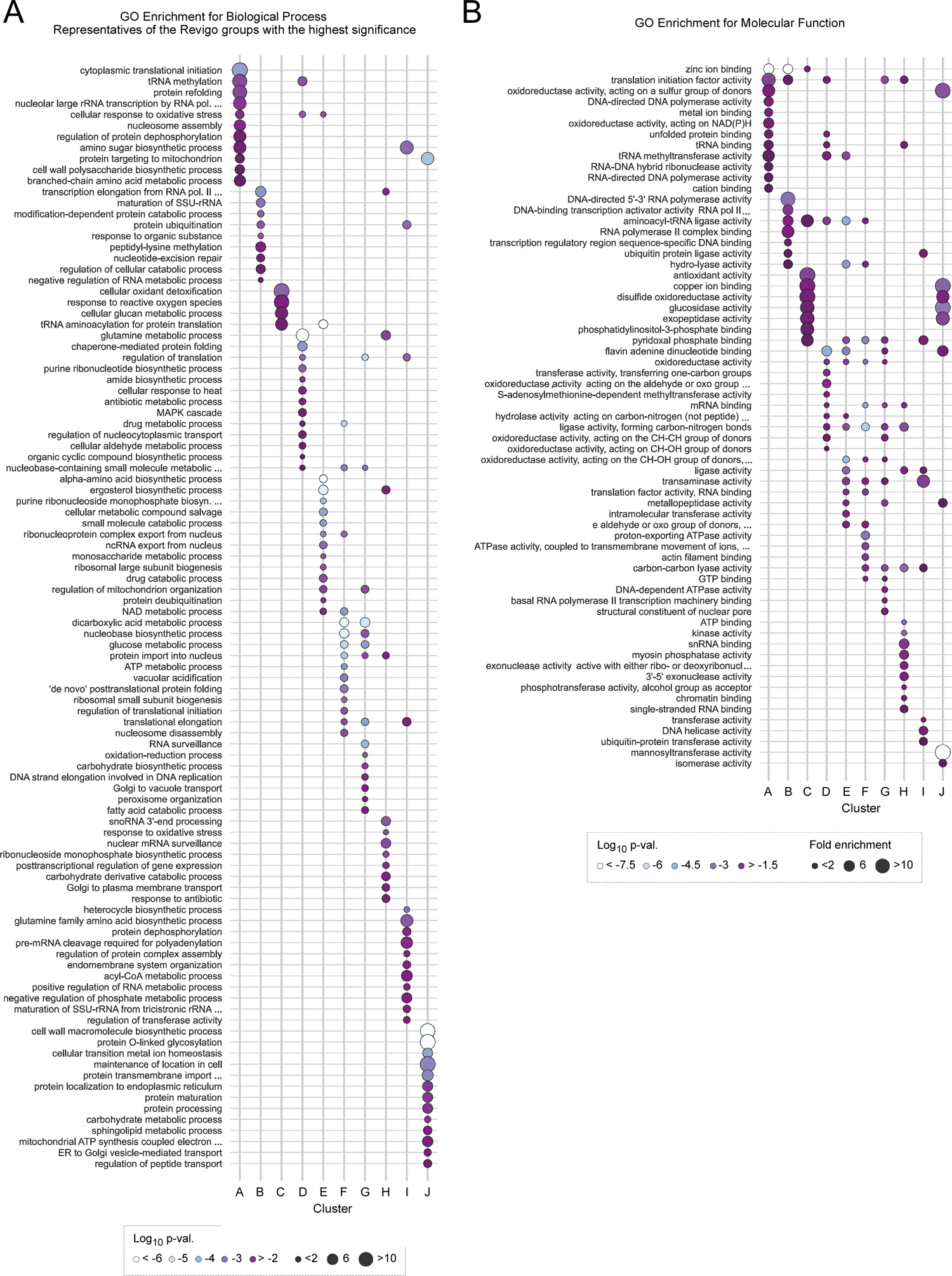
GO enrichment analysis for biological process and molecular function for individual clusters. **(A)** GO term enrichment analysis for biological process (BP) for individual clusters A-J. GO BP terms were grouped by Revigo based on the relation between GO terms for simplification of the graph. GO BP terms shown are representative of the Revigo groups with the highest statistical significance (Fisher’s test, Benjamini-Hochberg correction). A maximum of 15 best-scored terms are shown. Dot size, fold enrichment; dot colour, log_10_ uncorrected *p*-value. **(B)** GO term enrichment analysis for molecular function (MF) for individual clusters A-J. A maximum of 15 best-scored terms based on the highest statistical significance (Fisher’s test, Benjamini-Hochberg correction) are shown. Dot size, fold enrichment; dot colour, log_10_ uncorrected *p*-value.

**Supplementary Figure 4. (Related to Figure 5).**
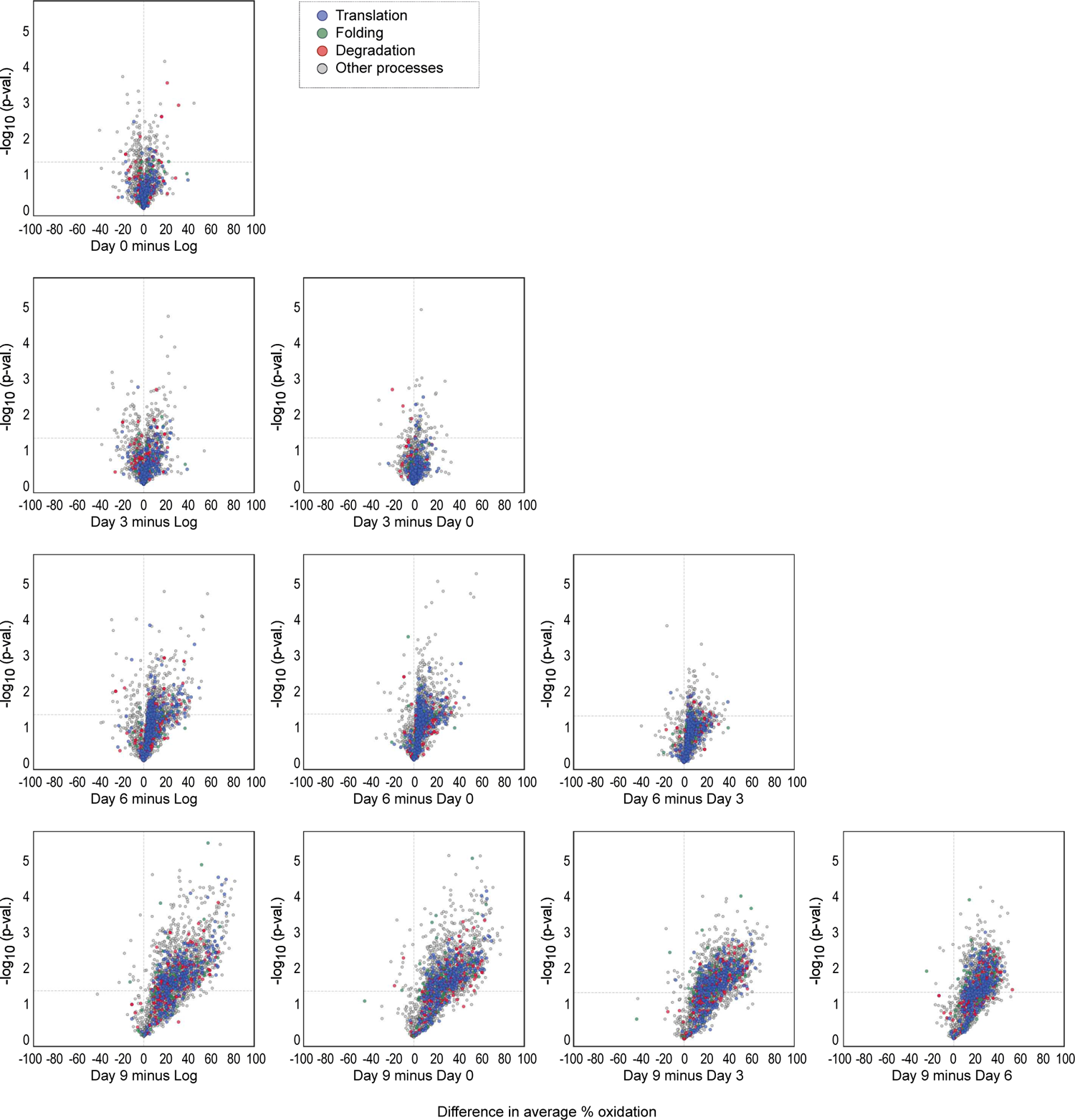
Pairwise comparison of oxidation of cysteine residues of proteostasis-regulating proteins during ageing. Pairwise comparison of *in vivo* cysteine modification during proliferation (Log) and different stages of chronological ageing (days 0, 3, 6, and 9) from the “yeast OxiAge” dataset (imputed values) for proteins involved in proteostasis regulation. Difference in % oxidation between time points is shown on the x-axis; -log_10_ *p*-value (two-tailed Welch’s *t*-test) is shown on the y-axis. Grey vertical line, difference = 0; grey horizontal line, p-value = 0.05. Grey dots, peptides of proteins that were not classified to proteostasis processes; dark blue dots, peptides of proteins that were classified into the regulation of cytoplasmic translation; green dots, peptides of proteins that were classified into the regulation of protein folding; light red dots, peptides of proteins that were classified into the regulation of protein degradation.

**Supplementary Figure 5. (Related to Figure 5).**
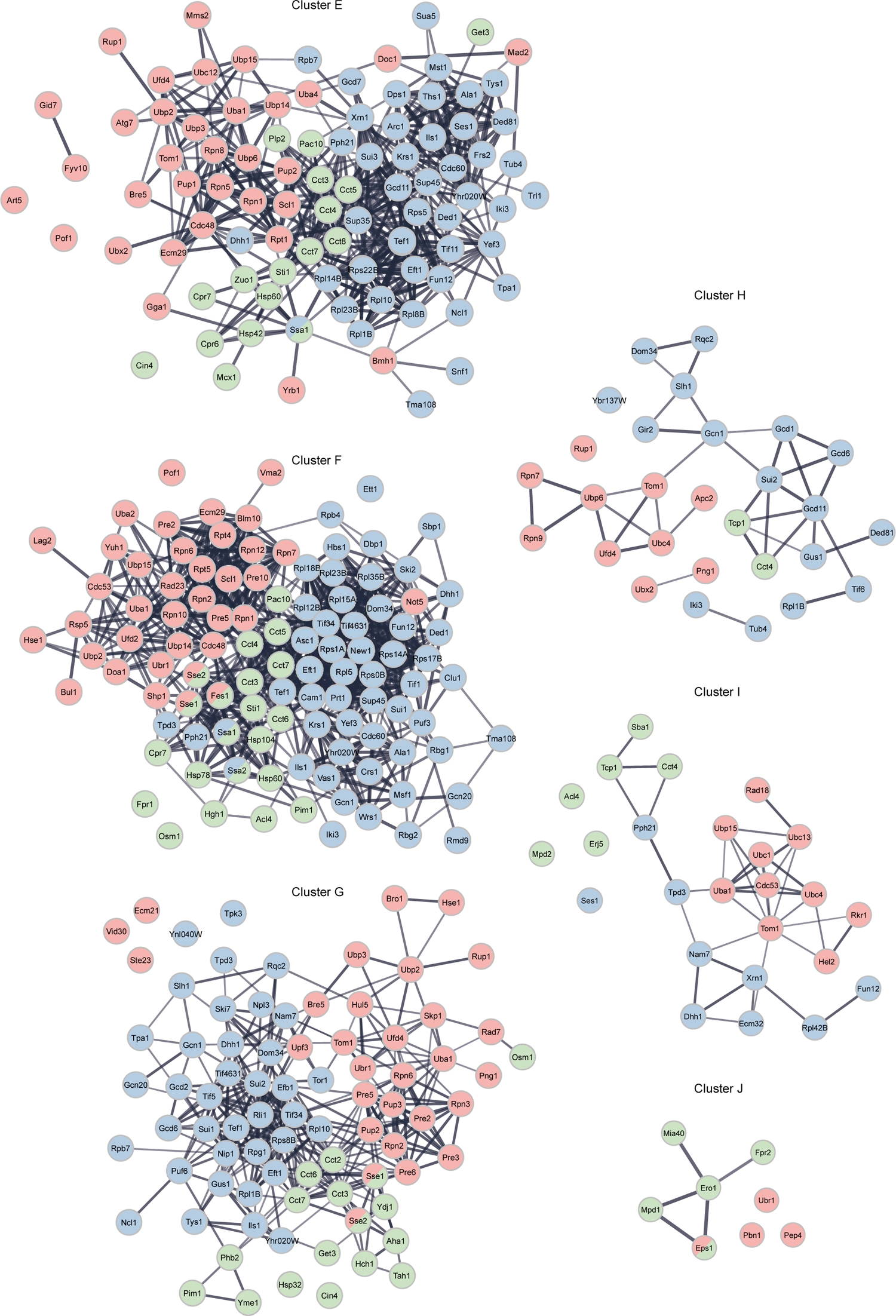
Protein association networks for proteostasis-regulating proteins within clusters E-J. STRING protein association network for proteostasis-regulating proteins for individual clusters E-J. Nodes, proteins; lines, interaction evidence. Line thickness, the strength of the evidence. Nodes are coloured based on the biological process the protein is involved in. Proteins involved in more than one process have more than one colour.

**Supplementary Figure 6. (Related to Figure 6).**
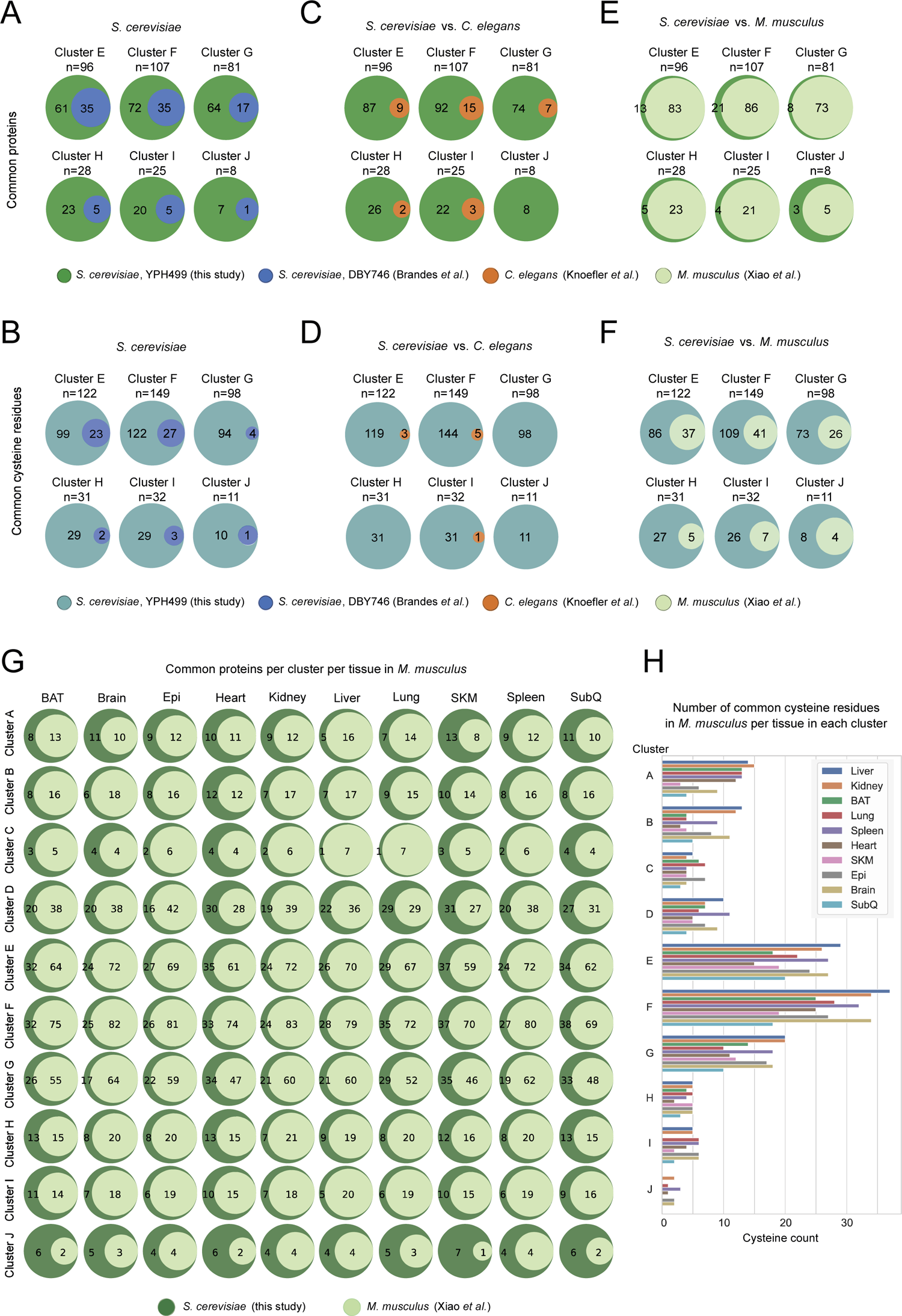
Cross-species comparison of oxidation of proteostasis-regulating proteins during ageing. (A-F) Pairwise comparison of commonly oxidized proteins (A, C, E) and cysteine residues (B, D, F) between species. Orthologs were compared to each cluster E-J individually. Green, unique proteins from “yeast OxiAge” dataset (A, C, E); turquoise, unique cysteine residues from “yeast OxiAge” dataset (B, D, F); blue, unique proteins or cysteine residues from yeast Brandes *et al*. dataset; brown, unique proteins or cysteine residues from worm Knoefler *et al*. dataset; pale green, unique proteins or cysteine residues from mice “Oximouse” Xiao *et al*. dataset. (G) Pairwise comparison of commonly oxidized proteins between the “yeast OxiAge” dataset and “Oximouse” (Xiao *et al*.) for individual clusters and mouse tissues. Green, unique proteins from the “yeast OxiAge” dataset; pale green, unique proteins from the “Oximouse” dataset. *BAT*, brown adipose tissue; *Epi*, epididymal fat; *SLM*, skeletal muscle; *SubQ*, subcutaneous fat. (H) Number of common cysteine residues between the “yeast OxiAge” and the “Oximouse” datasets per each mice tissue per cluster. *BAT*, brown adipose tissue; *Epi*, epididymal fat; *SLM*, skeletal muscle; *SubQ*, subcutaneous fat.

**Supplementary Figure 7. (Related to Figure 7).**
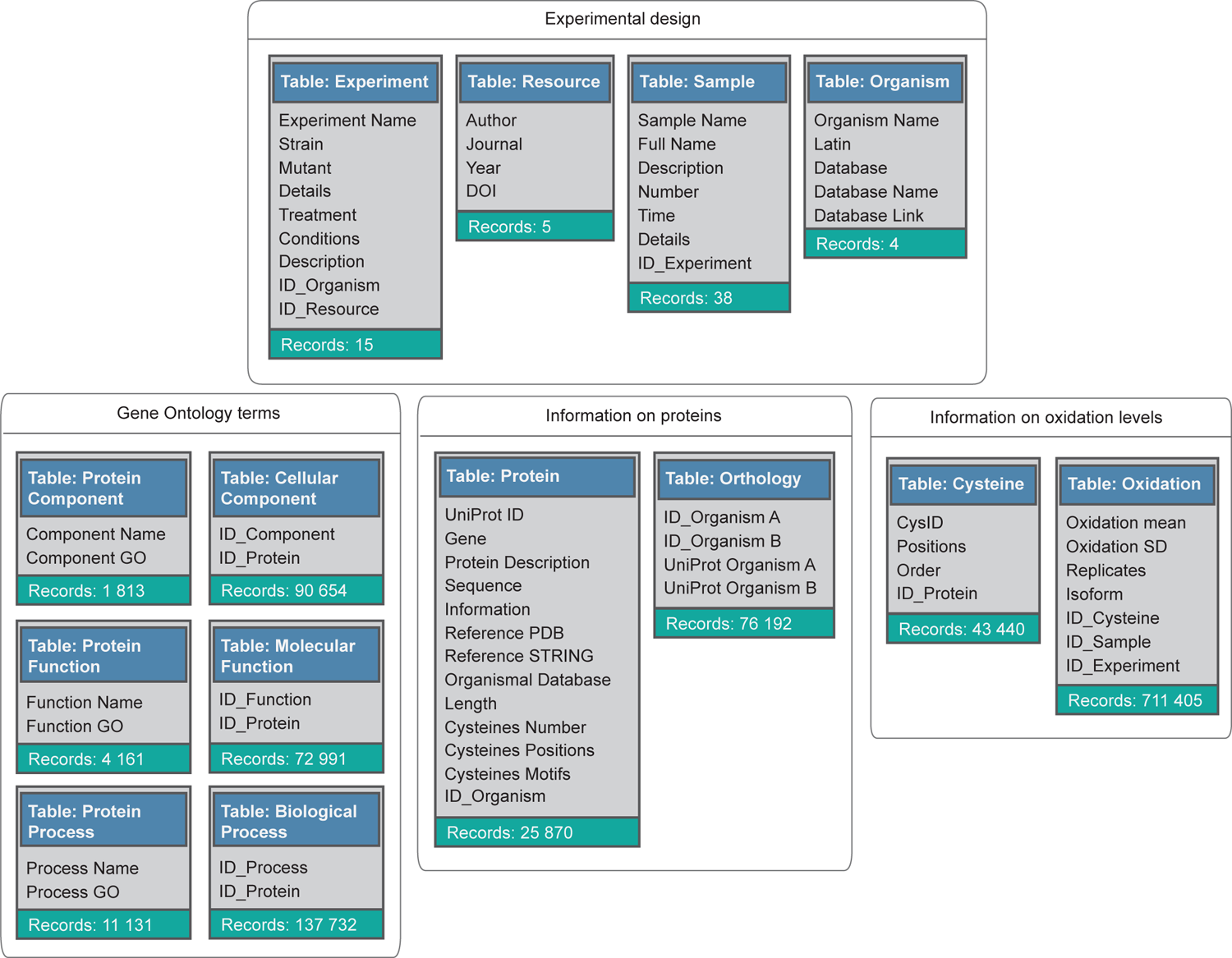
Design of the OxiAge database. (A) Design of the MySQL database containing information about the average % oxidation of aged proteome of *S. cerevisiae* (this study and Brandes *et al*.), *C. elegans* (Knoefler *et al*.), *D. melanogaster* (Menger *et al*.), and *M. musculus* (Xiao *et al*.). Unfiltered data was used to allow the user to perform custom filtering based on individual criteria. The database consists of 14 main tables divided into four groups (grey squares) based on function. Schemes of the tables contain the table name (blue background), the number of records (rows) within a table (sea-green background), and column names within each table. Column names starting with “ID” indicate the identification number referencing a record in another table.

**Supplementary Table 1.** The source file of mass spectrometry data comparing all detected peptides containing redox-sensitive thiols at different time points of chronological ageing in wild-type yeast *S. cerevisiae* (strain: YPH499).

**Supplementary Table 2.** The file of filtered mass spectrometry data comparing quantified peptides containing redox-sensitive thiols at different time points of chronological ageing in wild-type yeast *S. cerevisiae*. We refer to this data as “yeast OxiAge”.

**Supplementary Table 3.** Information about clusters A-J to which individual Cys-peptides from the “yeast OxiAge” dataset were assigned. Along with cluster names is information about amino acid sequences surrounding oxidized cysteine residue (motifs) and protein localization.

**Supplementary Table 4.** Gene ontology (GO) enrichment for biological process (BP) and molecular function (MF) of proteins assigned to clusters A-J in the “yeast OxiAge” dataset.

**Supplementary Table 5.** List of proteins from the “yeast OxiAge” dataset assigned to one of three main proteostasis-regulating processes: cytoplasmic translation, protein folding or protein degradation.

**Supplementary Table 6.** List of filtered peptides from *S. cerevisiae* Brandes *et al*. dataset, *C. elegans* Knoefler *et al*. dataset, *D. melanogaster* Menger *et al*. dataset, and *M. musculus* Xiao *et al*. dataset.

**Supplementary Table 7.** List of all proteins commonly oxidized during ageing between four species and five datasets: yeast *S. cerevisiae*, strain YPH499 from this study (“yeast OxiAge”), yeast *S. cerevisiae*, strain DBY746 from Brandes *et al*. dataset, nematode *C. elegans* from Knoefler *et al*. dataset, *D. melanogaster* from Menger *et al*. dataset, and mouse *M. musculus* from Xiao *et al*. dataset.

**Supplementary Table 8.** List of proteins commonly oxidized during ageing for the “yeast OxiAge” proteostasis sub-dataset and Brandes *et al*., Knoefler *et al*., Menger *et al*., and Xiao *et al*. datasets, pairwise comparison.

**Supplementary Table 9.** List of cysteine residues commonly oxidized during ageing for the “yeast OxiAge” proteostasis sub-dataset and Brandes *et al*., Knoefler *et al*., Menger *et al*., and Xiao *et al*. datasets, pairwise comparison.

